# Distributing Aminophospholipids Asymmetrically Across Leaflets Causes Anomalous Membrane Stiffening

**DOI:** 10.1101/2022.12.20.521165

**Authors:** Moritz PK Frewein, Paulina Piller, Enrico F Semeraro, Orsolya Czakkel, Yuri Gerelli, Lionel Porcar, Georg Pabst

## Abstract

We studied the mechanical leaflet coupling of prototypic mammalian plasma membranes using neutron spin-echo spectroscopy. In particular, we examined a series of asymmetric phospholipid vesicles with phosphatidylcholine and sphingomyelin enriched in the outer leaflet and inner leaflets composed of phosphatidylethanolamine / phosphatidylserine mixtures. The bending rigidities of most asymmetric membranes were anomalously high, exceeding even those of symmetric membranes formed from their cognate leaflets. Only asymmetric vesicles with outer leaflets enriched in sphingolipid displayed bending rigidities in conformity with these symmetric controls. We performed complementary small-angle neutron and X-ray experiments on the same vesicles to examine possible links to structural coupling mechanisms, which would show up in corresponding changes in membrane thickness. Additionally, we estimated differential stress between leaflets originating either from a mismatch of their lateral areas or spontaneous curvatures. However, no correlation with asymmetry-induced membrane stiffening was observed. To reconcile our findings, we speculate that an asymmetric distribution charged or H-bond forming lipids may induce an intraleaflet coupling which increases the weight of hard undulatory modes of membrane fluctuations and hence the overall membrane stiffness.

**SIGNIFICANCE**
Biological membranes not only act as the outer barriers of cells and organelles, but host a delicate machinery of proteins, which is sensitive not only to the surrounding lipid composition, but also to the mechanical properties of the bilayer. Little is known about the effect of transbilayer lipid asymmetry, a hallmark of all plasma membranes, on bilayer structure and dynamics. The results of this study indicate that the composition of the plasma membrane leads to unique mechanical coupling mechanisms, suggesting that the regulation of lipid asymmetry through flipases/flopases and scramblases could be an important means to tune the mechanical stiffness of the membrane.

## INTRODUCTION

Cellular envelopes contain a large number of lipid species that are distributed asymmetrically between the two leaflets of the bilayer. For example, mammalian plasma membranes are known to be composed of an outer leaflet enriched in cholinephospholipids, while the majority of the aminophospholipids are confined to the inner leaflet (1, 2). Membrane asymmetry is generated and maintained by an array of enzymes termed flipases, flopases, and scramblases (3). However, the physiological need or benefits for membrane asymmetry are far from being understood. For example, the different chemical potentials stored in both leaflets might induce a coupling that can be exploited, e.g. in signaling or transport processes such as endo/exocytosis even in the absence of protein (4, 5). Currently conceived lipid-mediated coupling mechanisms consider either intrinsic lipid curvature, headgroup electrostatics, cholesterol flip-flop, dynamic chain interdigitation, and differential stress between lipid leaflets (6–8).

Transleaflet coupling has been reported to induce domains in non-domain forming lipid mixtures (9, 10), affect lateral lipid diffusion (11), or cross-link the melting transitions of individual leaflets (7). Recently, we reported that the extent of hydrocarbon interdigitation allows to tweak the lateral packing of lipids in the apposing leaflet through competing cohesive and entropic interactions of the acyl chains (12). Additionally, membrane asymmetry can lead to distinct effects on its elastic behavior, either due to lipid specific properties (e.g. size, shape), or simply lipid over/under-crowding of a given leaflet (8, 13). Reported experimental data for membrane asymmetry-induced stiffening agree at least qualitatively with this theory but are scarce (14–17). In particular, none of the previous reports interrogated the effect of asymmetrically distributed charged lipids on the bilayer’s bending rigidity. Note that charged lipids are able to increase the stiffness of symmetric bilayers significantly (18, 19).

This prompted us to perform bending rigidity measurements on asymmetric membranes with — compared to previous reports — higher compositional complexity. In particular, we engineered mimics of mammalian plasma membranes with lipid architectures close to the reported distributions of phosphatidylcholine, sphingomyelin, phosphatidylethanolamine and phosphatidylserine in mammalian plasma membranes (2). Asymmetric large unilamellar vesicles (aLUVs) were produced using cylcodextrin-mediated lipid exchange (20) and studied by small-angle neutron and X-ray scattering (SANS, SAXS) and neutron spin-echo spectroscopy (NSE). Cholesterol was deliberately excluded from the present study, despite its abundance in mammalian plasma membranes. Cholesterol is known to induce lipid domains in lipid mixtures mimicking mammalian membranes (9, 10), which challenges nanostructural interpretation as attempted in this report. Leaving out cholesterol thus allowed us to probe pure phospholipid interactions without any interference from heterogeneities (domains) induced by cholesterol-lipid interactions.

We find that aLUVs with inner leaflets enriched either in palmitoyl oleoyl phosphatidylethanolamine (POPE), or POPE/phosphatidylserine (POPS) and outer leaflets containing palmitoyl oleoyl phosphatidylcholine (POPC), milk sphingomyelin (MSM), or equimolar mixtures of both lipids are always more rigid than symmetric vesicles of the same lipid composition. Moreover, most aLUVs were even more rigid than symmetric vesicles of their cognate leaflets, i.e. displayed anomalous stiffening. No anomalous stiffening was observed when only MSM was enriched in the outer leaflet, which can, due to its highly asymmetric hydrocarbon chain composition, interdigitate into the inner leaflet (12). Nanostructural data of the same samples allowed us to seek agreement with existing theories which couple membrane thickness (21) and differential stress (8) to the elastic behaviour of membranes. However, we could not find a satisfactory agreement. We speculate that charge and hydrogen bonding mediated intraleaflet interactions between aminophospholipids lead to a dominance of short wavelength fluctuations (hard modes), when these lipids are distributed asymmetrically in bilayers and hence to much increased bending rigidities. The absence of anomalous stiffening for MSM containing asymmetric membranes suggests that hydrocarbon chain interdigitation can alleviate this effect.

## MATERIALS AND METHODS

### Lipids, Chemicals and Sample Preparation

POPC, POPE, POPS, MSM, as well as egg sphingomyelin (ESM), dipalmitoylphosphatidylcholine (DPPC), dipalmitoylphosphatidylglycerol (DPPG), dioleoyl phosphatidylcholine (DOPC), and palmitoyl oleoyl phosphatidylglycerol (POPG) were purchased from Avanti Polar Lipids (Alabaster, AL, USA) and used without further purification. Chloroform, methanol (*pro analysis* grade), sucrose and methyl-*β*-cyclodextrin (m*β*CD) were obtained from Merck KGaA, Darmstadt, Germany. We prepared asymmetric unilamellar vesicles following the heavy donor cyclodextrin exchange protocol (20). Acceptor and donor lipids were weighed (ratio 1:2 mol/mol), dissolved separately in a chloroform/methanol mixture (2:1, vol/vol) and dried under a soft argon stream in a glass vial. Acceptor vesicles were prepared from either DPPC with 5 mol% DPPG, POPE with 10 mol% POPG, or POPE with 30 mol% POPS. Donor vesicles consisted of ESM, MSM, POPC or equimolar POPC:MSM mixtures and did not contain any charged lipid.

The resulting dry lipid films were kept over night in vacuum to ensure the evaporation of all solvent and subsequently hydrated. Acceptor vesicles were prepared in ultrapure H_2_O containing 25 mM NaCl (lipid concentration: 10 mg/ml) by 1 h incubation at 50 °C with intermittent vortex mixing and 5 freeze/thaw cycles. Donor vesicles were hydrated analogously, but in a 20 wt% sucrose solution (lipid concentration: 20 mg/ml). Acceptor vesicles were extruded at 50 °C using a handheld mini extruder (Avanti Polar Lipids, AL, USA) with a 100 nm pore diameter polycarbonate filter 31 times or until reaching a polydispersity index < 10%. The latter parameter was monitored by dynamic light scattering (DLS) using a Zetasizer NANO ZS90, (Malvern Panalytical, Malvern, UK) ensuring the presence of exclusively large unilamellar vesicles (LUVs). Multilamellar vesicles (MLVs) of donor lipids were diluted 20-fold with water and centrifuged at 20 000 g for 30 min. The supernatant was discarded. The resulting pellet was suspended in a 35 mM m*β*CD solution (lipid:m*β*CD 1:8 mol/mol) and incubated for 2 h at 50 °C, while being shaken at a frequency of 600 min−1. Acceptor vesicles were added and incubated for another 15 min. The exchange was stopped by diluting the mixture 8-fold with water and centrifuging at 20 000 g for 30 min. The supernatant containing the asymmetric vesicles was then concentrated to < 500 *μ*l using 15 ml-Amicon centrifuge filters (Merck, 100 kDa cut-off) at 5000 g. To remove residual m*β*CD and sucrose, and to exchange H_2_O by D_2_O, filters were filled with D_2_O and re-concentrated in 3 cycles. The final vesicle sizes were again measured by DLS to ensure the absence of donor MLVs.

Scrambled LUVs were produced by drying asymmetric vesicles in a rotary evaporator at 30 °C and pressures around 50 Pa. The resulting films were thereafter rehydrated with D_2_O and extruded as described above. Symmetric reference LUVs for inner and outer leaflets were prepared in pure D_2_O analogously. Inner leaflet reference LUVs contained 90 mol% acceptor lipid and 10 mol% donor lipid, while outer leaflet reference LUVs were mixtures of 30 mol% acceptor and 70 mol% donor lipid.

### Compositional Analysis using Gas Chromatography (GC) and High Performance Thin Layer Chromatography (HPTLC)

The overall vesicle composition of all aLUVs — except for POPE^in^ /POPC^out^ and (POPE/POPS)^in^ /POPC^out^ — was determined by characterizing the fatty acid composition via GC. For this procedure, fatty acid methyl esters (FAMEs) were prepared upon incubation with a methanolic-H_2_SO_4_ solvent mixture. GC measurements were done using a GC 2010 Plus (Shimadzu), equipped with a split/splitless injector and a SGE BPX70-Cyanopropyl Polysilphenylene-siloxane column (25 m by 0.22 mm ID and 0.25 *μ*m film thickness) as described in (22). For calibration, we used logarithmically spaced concentration series of DPPC, DOPC, ESM (mostly 16:0 acyl chains), and MSM (mostly 22:0, 23:0 and 24:0 acyl chains). Lipid composition was given by interpolating through these calibration points.

POPE^in^ /POPC^out^ and (POPE/POPS)^in^ /POPC^out^ aLUVs were analyzed by HPTLC. After lipid extraction against organic solvent — CHCl_3_/MeOH 2:1 (vol/vol) — samples were spotted on silica plates (Sigma-Aldrich, Steinheim, Germany) with the automatic TLC sampler 4 (CAMAG, Switzerland). The mobile phase in the developing chamber was a solvent mixture composed of 32.5:12.5:2 (vol/vol/vol) CHCl_3_/MeOH/H_2_O. After drying, the plate was immersed in a developing bath (5.08 g MnCl_2_ dissolved in 480 ml H_2_O, 480 ml EtOH and 32 ml H_2_SO_4_), which is sensitive to double bonds, and dried for 15 min at 120 °C. To quantify the lipid concentrations the plate was scanned with the TLC scanner 3 (CAMAG, Switzerland) and further analyzed with WinCats software.

### Measuring Asymmetry via Nuclear Magnetic Resonance (NMR) Spectroscopy

We used ^1^H-NMR spectroscopy to determine the ratio of donor lipids in inner and outer leaflets by exposing the aLUVs to the shift reagent Pr^3+^ as detailed previously (20). Choline, which is present in all headgroups of here used donor lipids, gives a strong ^1^H-NMR signal, which is shifted if the headgroup is surrounded by Pr^3+^. Adding the reagent after sample preparation affects only outer leaflet headgroups within the timescale of the experiment and thereby allows to discriminate against the signal of inner leaflet cholinelipids. Experiments were performed on a Avance III 600 MHz spectrometer (Bruker, Billerica, MA) using the Bruker TopSpin acquisition software. A ^1^H pulse sequence including water suppression (23) was used to collect 64 transients at 50 °C. Data were processed with TopSpin 3.6 applying a line-broadening parameter of 1.5 Hz. In contrast to former measurements, we did not shift the outer leaflet peak to a certain position, but added enough Pr^3+^ to decrease its intensity far below those of its neighboring peaks (4 *μ*l of a 20 mM Pr^3+^/D_2_O solution added to 500 *μ*l lipid vesicle solution of about 0.5 mg/ml). The inside/outside ratio of donor lipid is given by the areas of the peak of the total cholines before and after adding the shift reagent; see the supplementary information (SI) for more information.

### Small-Angle Neutron & X-ray Scattering (SANS, SAXS)

SANS measurements were performed at D22, Institut Laue-Langevin, Grenoble, France and are available from DOI (10.5291/ILL-DATA.9-13-953, 10.5291/ILL-DATA.9-13-997). We used a multidetector setup with two ^3^H-detectors, one of which was positioned at sample detector distances SDD = 5.6 m, or 17.8 m with collimations of 5.6 m and 17.8 m, respectively, while the second detector was positioned out of center of the direct neutron beam at SDD = 1.3 m. The neutron wavelength was 6 Å (Δλ /λ = 10%). Samples were filled in Hellma 120-QS cuvettes of 1 mm pathway and heated to 50 °C using a water bath circulator. Lipid concentrations were about 5 mg/ml in 100 % D_2_O. Data reduction, including flat field, solid angle, dead time and transmission corrections, intensity normalization by incident flux and subtraction of contributions from empty cell and solvent were performed using GRASP (ILL).

SAXS data were recorded at BM29, ESRF, Grenoble, France, using a photon energy of 15 keV and a Pilatus3 2M detector at SDD = 2.867 m (data available at DOI: 10.15151/ESRF-ES-514136943). Samples were measured at a concentration of 10 mg/ml, at 50 °C and exposed for 40 s (20 frames of 2 s each) in a flow-through quartz glass capillary (diameter: 1 mm). Data reduction and normalization was done by the automated ExiSAXS system, SAXSutilities 2 (ESRF) was used for subtracting contributions from solvent and glass capillary.

SANS and SAXS data were analyzed jointly in terms of Gaussian-type volume distribution functions of quasi-molecular lipid fragments across the bilayer, using our recently reported advanced scattering density model for aLUVs (12) (for details, see the SI). Compared to this previous study, we did not use lipids with deuterated hydrocarbons. Firstly, because our NSE experiments required fully protiated samples for optimal contrast, and secondly, because of their limited commercial availability. Consequently we are not able to discriminate the structure of the individual leaflets. Structural parameters, such as the lateral area per lipid, *A*, are thus averaged over both leaflets. The chain thickness *D*_*C*_ of each leaflet was estimated from the position of the terminal methyl group. Note that this might not necessarily correspond to the center at the membrane, especially for samples containing the highly chain asymmetric MSM (12).

Data were fitted following (24), but additionally constraining lipid composition of inner and outer leaflets with results from GC, HPTLC and ^1^H-NMR experiments as described by Eicher et al. (25). Further, we fixed the distance between the hydrophobic interface and the backbone *d*_BB_ = 0.9 Å, the width of the Gaussians describing backbone *σ*_BB_ = 2.1 Å and the outermost headgroup (choline/amine) *σ*_Chol_ = 3 Å and the smearing of the error-function describing the hydrophobic-hydrophilic interface *σ*_CH2_ = 2.5 Å. Volumes of the individual moieties of the lipids were taken from (26). In case of lipid mixtures, these values where calculated using molecular averaging and the experimentally determined estimates of leaflet composition (see results).

### Neutron Spin-Echo Spectroscopy (NSE)

NSE measurements were performed at the IN15 spin-echo spectrometer, Institut Laue-Langevin, Grenoble, France (data available at DOI: 10.5291/ILL-DATA.TEST-3125, 10.5291/ILL-DATA.9-13-953, 10.5291/ILL-DATA.9-13-997) (27). We recorded NSE data in the range of *q* = 0.020 Å^−1^to 0.111 Å^−1^ using λ = 8, 10 and 12 Å. Lipid dispersions (concentration ∼ 15 mg/ml) were filled into quartz-cells of 1 mm pathlength. The accessible Fourier times ranged, depending on wavelength, from 0.01 ns up to 300 ns. All experiments were performed at 50°C.

NSE data were analyzed in terms of the Zilman-Granek (ZG) theory (28) following the approach of Gupta et al. (29). In brief, the mean squared displacement (MSD), ⟨Δ*r*(*t*)^2^⟩, of the membrane from a flat geometry is related to the intermediate scattering function *S*(*q, t*) assuming a Gaussian distribution of fluctuations, which at the same time is required to follow the ZG model

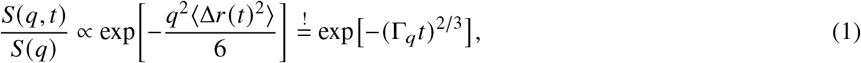

where *S*(*q)* = *S*(*q*, 0) is the elastic structure factor. The decay constant Γ_*q*_ is connected to the ‘effective’ or ‘dynamic’ bending rigidity 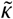.

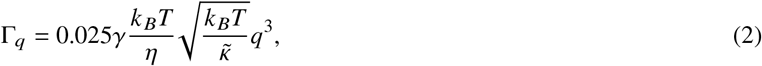

where *γ* ≈ 1 for 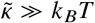 (28), *η* is the solvent viscosity 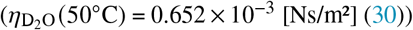 (30)), *k* _*B*_ is Boltzmann’s constant and *T* is the absolute temperature. The advantage of the MSD approach is the ability to discriminate against deviations from the ZG prediction for the decay of *S*(*q, t*)/*S*(*q*) either due to non-Gaussian fluctuations occurring at short Fourier times, or due to vesicle diffusion dynamics, which dominate current data at long Fourier times (29). This approach avoids any ‘artificial’ scaling by particle diffusion or interfacial viscosity (supplementary Fig. S3), which has led to significant controversies about bending rigidity values *κ* on absolute scale reported from NSE (31).

After ensuring the proper Fourier time range for the application of the ZG model we derived *κ* following the model of Watson and Brown (32). This theory takes internal dissipation into account and arrives at 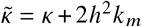, where *h* is the position of the neutral surface and *k*_*m*_ is the monolayer area compressibility modulus. Further, using 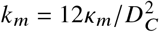, as predicted by the polymer brush model (21), and *κ*_*m*_ = *κ*/2 (33) for each leaflet, one arrives at

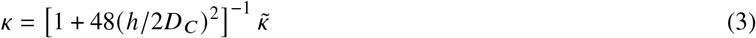

The exact position of *h* is not entirely clear and *h*/2*D*_*C*_ and different groups report values between 0.25 and 0.6 (34). Following our work on intrinsic lipid curvatures (35) we assume that the neutral surface coincides with the lipid’s backbone and calculate *h*/(2*D*_*C*_) from results of our SAXS/SANS SDP-analysis.

## RESULTS

### Transbilayer Asymmetry and Structure

We prepared a set of seven differently composed aLUVs using cyclodextrin-mediated exchange. Two different acceptor vesicles, POPE and POPE/POPS (7:3 mol/mol), were prepared. The latter mixture mimics to first order the natural composition of the inner leaflet of mammalian plasma membranes (2). Exchange was performed with four different donor MLVs: MSM, ESM, POPC and POPC/MSM (1:1 mol/mol), the latter of which mimics the outer leaflet of plasma membranes. Further, comparing MSM and ESM allowed us to probe the effects of chain length distribution in sphingolipids. The achieved total exchange and asymmetries are detailed in Tab. 1. Overall, exchange of the outer leaflet showed significant variations and was highest for POPE^in^/POPC^out^, where 65 mol% of acceptor lipid was replaced, a value which agrees well with a previous report (7). Exchange was in general low for both sphingolipids (< 30 mol%). This might also affect the molecular ratio of aLUVs containing POPC/MSM mixtures, i.e. the final POPC/MSM molar rations might be < 1. Note that ^1^H-NMR experiments performed on the chosen lipid moieties cannot be used to differentiate between phosphatidylcholines and sphingolipids. Such an analysis would require the application of headgroup deuterated species (36), which is beyond the scope of the present study.

In the next step, we determined the corresponding membrane structural parameters using the joint analysis of SANS and SAXS data. The transbilayer structure was parsed into the volume distributions of phosphate (PCN), choline-CH_3_ (CholCH3), glycerol or sphingosine backbone (BB), methylene/methine (chains), and methyl (CH3) groups, including also bound water in the headgroup region (12). The applied model is in principle capable of deriving leaflet-specific structural details (12). However, the structural resolution of the individual leaflets is poor in the absence of deuterated phospholipids. Deriving leaflet specific structural details is specifically challenged by the fact that the location of the terminal methyl groups — especially for the highly chain asymmetric MSM — might not be exactly at the center of an asymmetric bilayer (12, 24). Therefore, we report the areas per lipid averaged over both leaflets, *A*_av_, for aLUVs in Table 1; non-averaged, leaflet-specific structural data are listed in Tab. S3; fits are shown in Fig. S5. Additionally, we report in Table 1 the corresponding structural parameters for scrambled vesicles, that is LUVs with the same, but symmetric lipid composition; fits to SANS/SAXS data are shown in Figs. S10 – S16 and structural details in Tabs. S5 – S7. We note deviations in the SAXS data fits of some samples at very low scattering angles. The origin of these deviations is most likely due to issues with assigning proper contrast in the headgroup regime. However, because membrane structure dominates scattering at higher *q*, the impact of this mismatch at low *q* on the derived structural parameters is rather low; see also (24) for more details.

**Table 1:**
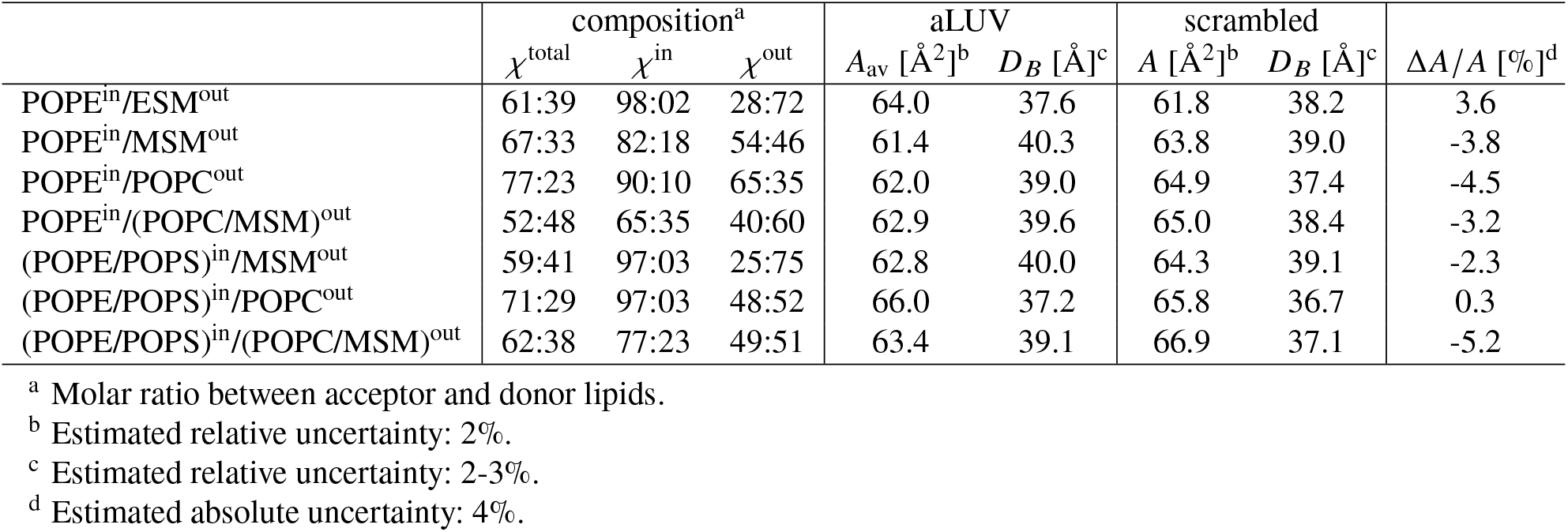
Composition and structure of presently studied aLUVs and scrambled LUVs.

| | composition<sup>a</sup> | | | aLUV | | scrambled | | $\Delta A/A$ [%]<sup>d</sup> |
| --- | --- | --- | --- | --- | --- | --- | --- | --- |
| | $\chi^{\text{total}}$ | $\chi^{\text{in}}$ | $\chi^{\text{out}}$ | $A_{\text{av}}$ [$\text{\AA}^2$]<sup>b</sup> | $D_B$ [$\text{\AA}$]<sup>c</sup> | $A$ [$\text{\AA}^2$]<sup>b</sup> | $D_B$ [$\text{\AA}$]<sup>c</sup> | |
| POPE<sup>in</sup>/ESM<sup>out</sup> | 61:39 | 98:02 | 28:72 | 64.0 | 37.6 | 61.8 | 38.2 | 3.6 |
| POPE<sup>in</sup>/MSM<sup>out</sup> | 67:33 | 82:18 | 54:46 | 61.4 | 40.3 | 63.8 | 39.0 | -3.8 |
| POPE<sup>in</sup>/POPC<sup>out</sup> | 77:23 | 90:10 | 65:35 | 62.0 | 39.0 | 64.9 | 37.4 | -4.5 |
| POPE<sup>in</sup>/(POPC/MSM)<sup>out</sup> | 52:48 | 65:35 | 40:60 | 62.9 | 39.6 | 65.0 | 38.4 | -3.2 |
| (POPE/POPS)<sup>in</sup>/MSM<sup>out</sup> | 59:41 | 97:03 | 25:75 | 62.8 | 40.0 | 64.3 | 39.1 | -2.3 |
| (POPE/POPS)<sup>in</sup>/POPC<sup>out</sup> | 71:29 | 97:03 | 48:52 | 66.0 | 37.2 | 65.8 | 36.7 | 0.3 |
| (POPE/POPS)<sup>in</sup>/(POPC/MSM)<sup>out</sup> | 62:38 | 77:23 | 49:51 | 63.4 | 39.1 | 66.9 | 37.1 | -5.2 |

<sup>a</sup> Molar ratio between acceptor and donor lipids.
<sup>b</sup> Estimated relative uncertainty: 2%.
<sup>c</sup> Estimated relative uncertainty: 2-3%.
<sup>d</sup> Estimated absolute uncertainty: 4%.

To facilitate comparison between aLUVs and scrambled LUVs we additionally report the relative change in lipid area 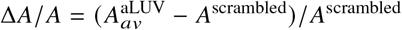 in Tab. 1. Overall, we observe that aLUVs are laterally more condensed than their scrambled analogs. Exceptions to this finding are POPE^in^ /ESM ^out^ with Δ*A*/*A* > 0 and (POPE/POPS)^in^ /POPC^out^ bilayers (Δ*A*/*A* ∼ 0).

### Bending Elasticity

NSE experiments were performed on the same exact samples studied by SANS/SAXS to avoid potential artifacts from sample history, e.g. sample to sample variations of lipid exchange efficiencies. The principles of the applied analysis are demonstrated in Fig. 2 for POPE^in^/POPC^out^ aLUVs. The MSD plot shows an overlap of data between *t* = 5 – 100 ns and *q*-values between ∼ 0.07 – 0.11 Å^−1^, all following a *t*^2/3^-slope as predicted by the ZG theory (28). Data that do not follow the ZG theory have been omitted from further analysis. Figure 2 also demonstrates the sensitivity of the technique to fluctuation differences between symmetric and asymmetric bilayers.

**Figure 1:**
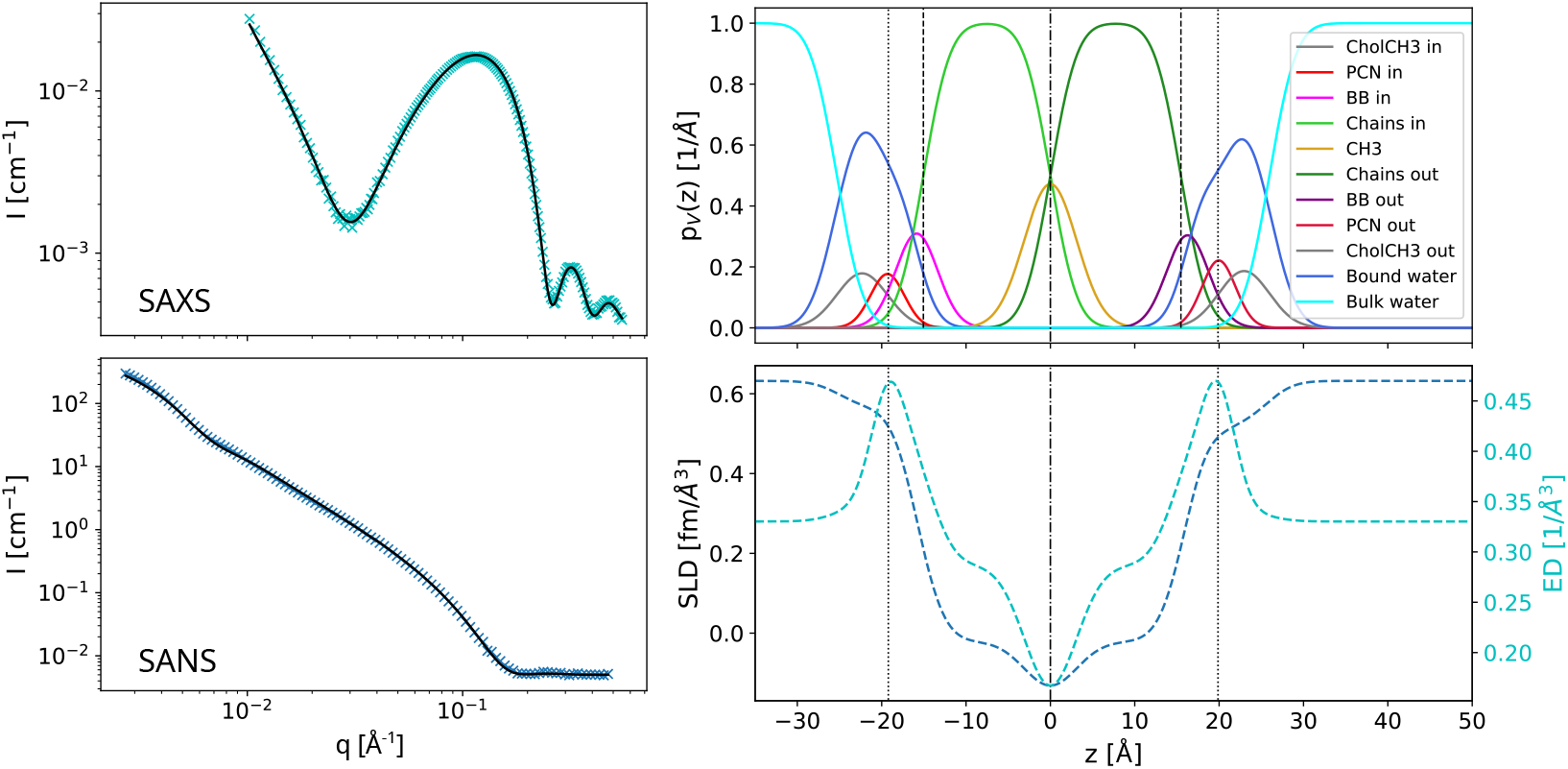
Structure of (POPE/POPS)^in^/(MSM/POPC)^out^ aLUVs as revealed by joint SAXS/SANS data analysis. Panels A and C show fits (black lines) to SAXS and SANS data (50 °C). Panel B gives the derived volume probability profiles for, and panel D the neutron scattering length density (SLD) and electron density (ED).

**Figure 2:**
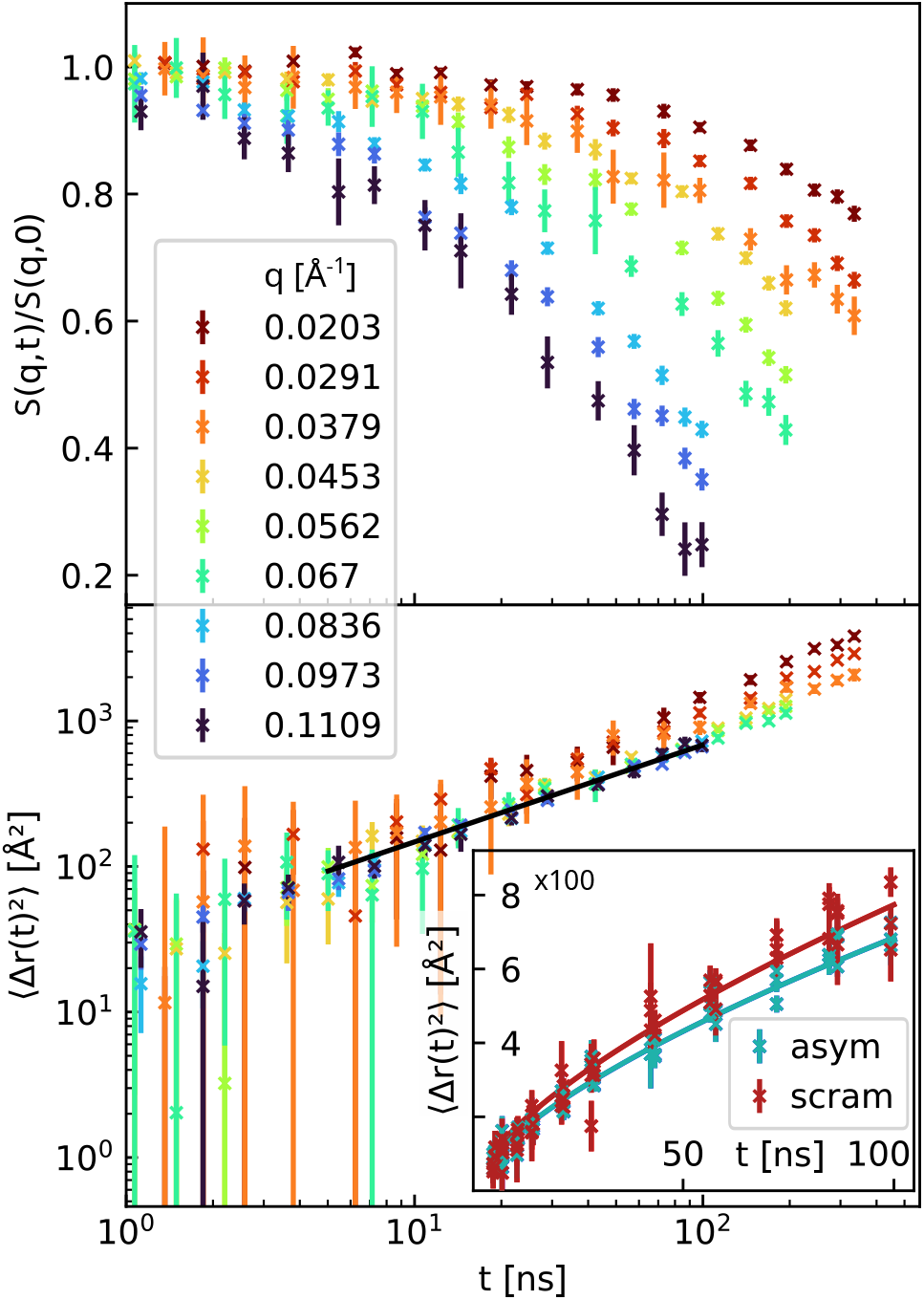
Principles of NSE data analysis. The upper panel shows raw NSE of POPE^in^/POPC^out^ aLUVs. Data below 5 ns was used to normalize *S*(*q, t*) /*S*(*q*, 0) and has been omitted in part in the plot due to experimental uncertainties. The lower panel presents the corresponding MSD (lower panel). The solid black line marks the range fitting the *t*^2/3^-slope predicted by the Zilman-Granek model, and which is used for the bending rigidity analysis. The insert compares the MSD data of the aLUVs to scrambled vesicles of the same lipid composition and demonstrates the sensitivity of the technique to differences in fluctuations dynamics.

We first tested our NSE analysis using DPPC LUVs at 50 °C as a benchmark system. Deriving the membrane bending rigidity requires, according to Eq. (3), the neutral plane, *h* and the hydrocarbon thickness *D*_*C*_. Assuming that *h* coincides with the position of the glycerol backbone (35), we arrive at *h*/(2*D*_*C*_) = 0.53 ± 0.07 using previously reported data for DPPC bilayers (12). This result is well within the range of previously reported values for *h*/(2*D*_*C*_) (34). Application of Eq. (3) then leads to *κ* = 12.0 ± 0.4 k_*B*_T, which is almost three times lower than the bending rigidity of DPPC reported from other techniques (37). Note that alternative NSE evaluations of the same data also do not yield satisfactory agreement with literature values (Fig. S3). It has been suggested that including a separate tilt modulus in the NSE data analysis should be considered to harmonize *κ*-values from NSE with other techniques (38). Indeed, Guler et al. (39) reported *κ* ∼ 15.0 k_*B*_T for DPPC — a value which is very close to our present result — using X-ray scattering, without correcting for the tilt modulus. Only later this value was corrected to *κ* ∼ 30 k_*B*_T by including the tilt modulus in the analysis (37). Revising the theory for NSE to bring *κ* to correct absolute scale levels is beyond the scope of the present study. Here it suffices to note that NSE is sensitive to relative changes of undulatory motions; see also inert to Fig. 2. We will thus discuss only the relative differences of *κ* between the studied samples.

### Symmetric References of Inner and Outer Leaflets

We performed additional NSE experiments on symmetric control samples composed either of POPC, MSM, ESM and POPE, or POPE/POPS (7:3 mol/mol) mixtures. These samples allowed us to estimate the bending rigidities of each leaflet for the aLUVs in the absence of any coupling mechanism (see below).

Table 2 summarizes the corresponding major results from SANS/SAXS and NSE experiments (see also Figs. S10 – S16, Tabs. S5 – S7). We briefly note some interesting elastic properties of these simple lipid systems: Firstly, the bending rigidity of POPC is lower than that of DPPC, despite its about equal membrane thickness (see the head-to-headgroup distances *D*_HH_, reported in Tab. S4). This is surprising in view of the well-accepted polymer brush model (21), which predicts 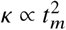, where *t*_*m*_ is the mechanical thickness of the membrane, which is frequently set equal to *D*_HH_. A recent computational approach showed that *t*_*m*_ can be smaller than *D*_HH_ for unsaturated hydrocarbons (40), which might explain our findings. Secondly, the bending rigidities of both sphingolipids are markedly higher than all other presently studied lipids. This indicates, additional contributions due to the different backbone-structure and the lipid’s ability to form intermolecular H-bonds. Thirdly, the addition of 30 mol% POPS to POPE increases the lateral area per lipid (Tab. S4), but does not lead to significant changes in *κ*. Possibly its concentration is still too low to cause membrane stiffening as reported for other charged bilayers (18, 19).

**Table 2:**
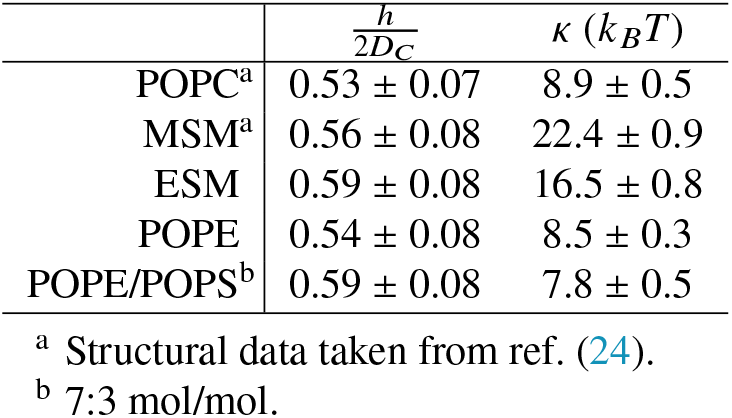
Structural and elastic properties of symmetric LUVs containing acceptor and donor lipids at 50 °C.

The bending rigidities of symmetric vesicles composed just of inner leaflet or outer leaflet lipid mixtures, detailed in Tab. 1, can now be estimated by interpolating the data reported in Tab. 2. Alternatively, one could have measured just vesicles with the exact leaflet compositions of Tab. 1. However, lipid exchange is difficult to predict. Thus, additional NSE, SAXS and SANS beamtimes would have been needed, just to measure the properties of these control samples. Our solution thus was to interpolate data, which was additionally aided by measuring the symmetric acceptor:donor lipid mixtures 9:1 (mol/mol) and 3:7 (mol/mol). These compositions roughly correspond to the inner and outer leaflet compositions of the aLUVs. The bending rigidity of POPC/MSM equimolar mixtures was calculated from the molecular average of *κ* of the individual lipids. Plotting *κ* of symmetric vesicles as a function of lipid composition, we observed clear linear trends for most studied bilayers (Fig. 3, Fig. S4), providing us with good confidence in the interpolated *κ*-values of the inner and outer leaflet mimics controls for comparing to the elasticity of aLUVs.

**Figure 3:**
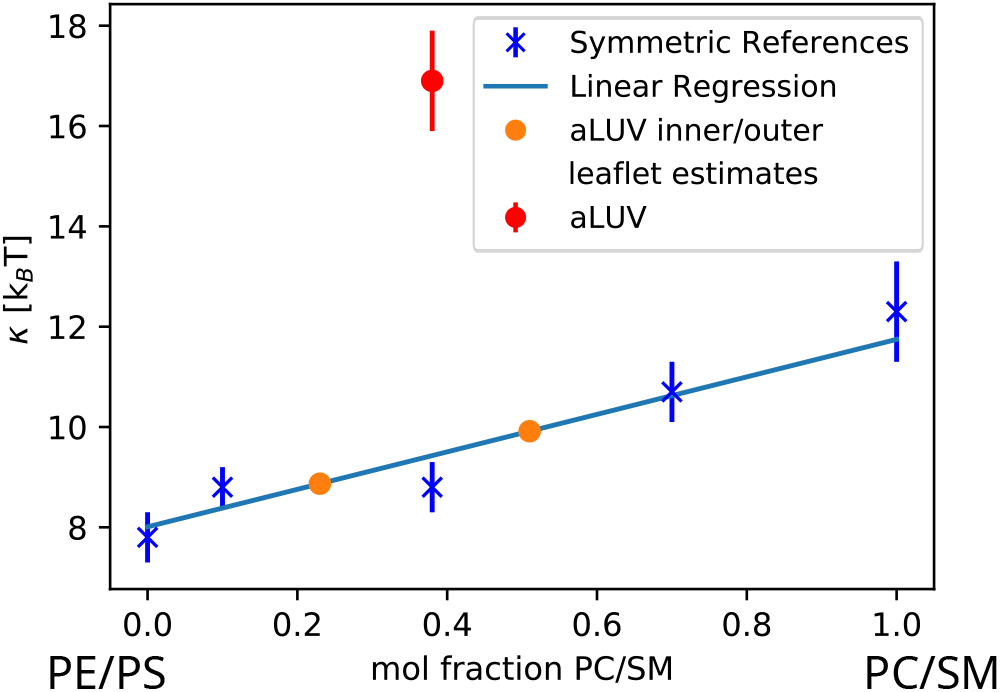
Variation of bending rigidity of symmetric POPE/POPS/POPC/MSM mixtures with POPC/MSM (1:1 mol/mol) concentration, abbreviated in the plot by PC/SM for clarity of presentation. A PC/SM mol fraction of 0 corresponds to POPE/POPS (7:3 mol/mol), here abbreviated by PE/PS. Orange circles represent *κ* estimates using the overall lipid compositions of Tab. 1. The red symbol shows *κ* of the corresponding aLUVs (see also Fig. 4.)

### Anomalous Elasticity of Asymmetric Bilayers

Figure 4 summarizes our results for *κ* of all studied aLUVs and compares them to the bending rigidities of three different symmetric LUVs: scrambled vesicles, and their cognate inner and outer leaflets. The bending rigidities of symmetric cognate leaflet samples correspond to upper and lower boundary expectation values for each system. In the case of uncoupled lipid leaflets *κ* of aLUVs should correspond to the arithmetic mean these boundary values (8). Measurements of scrambled LUVs provide an additional control for the effect of lipid distribution across the bilayer on bending fluctuations.

**Figure 4:**
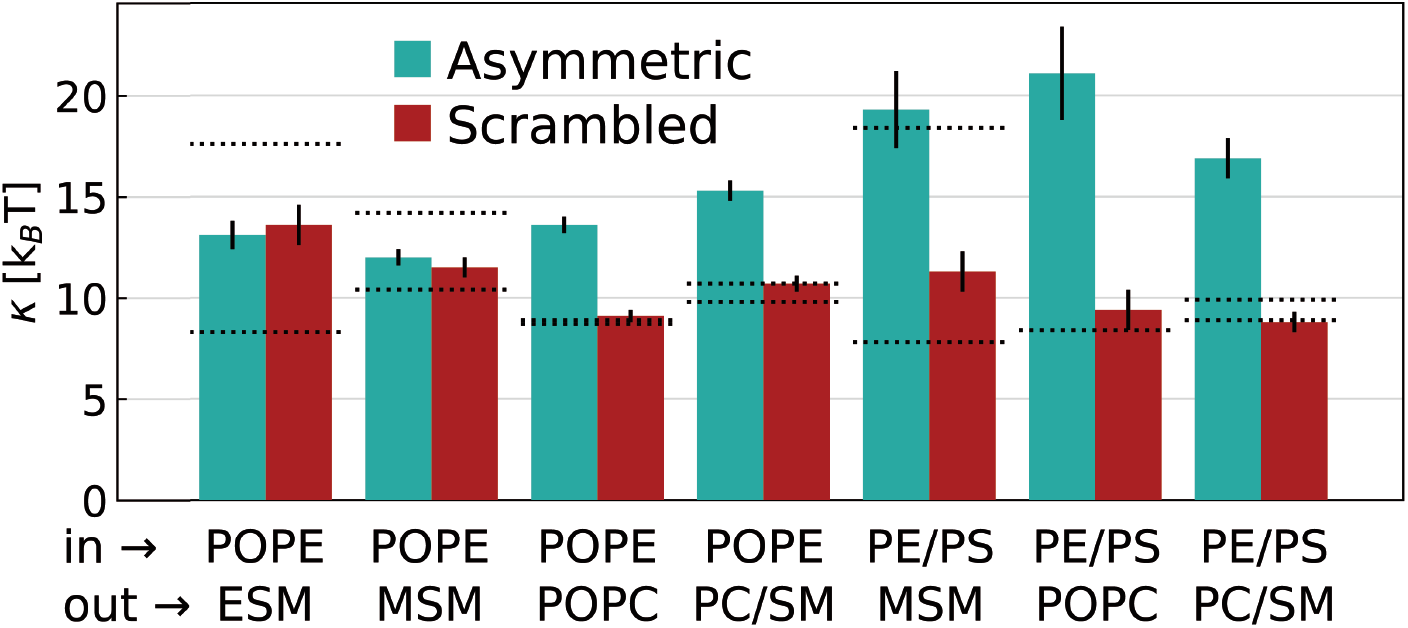
Bending rigidities *κ* measured by NSE for asymmetric and physically scrambled vesicles. Dotted lines correspond to the reference samples of inner and outer leaflet composition. The labels indicate the predominant lipids in inner and outer leaflet. The following mixtures have been abbreviated for clarity of display: PE/PS corresponds to POPE/POPS (7:3 mol/mol) and PC/SM to POPC/MSM (1:1 mol/mol).

Four aLUVs with inner leaflets enriched just in POPE were studied. These had outer leaflets with ESM, MSM, POPC, and equimolar mixtures of POPC and MSM. Differences in *κ* between asymmetric and scrambled membranes were negligible within experimental certainty for POPE/ESM and POPE/MSM vesicles. Further, bending rigidities of these asymmetric membranes corresponded to the arithmetic averages of their cognate leaflet vesicles, indicating the absence of trans-bilayer elastic coupling. This seems to contrast a recent NSE study on POPE^in^/ESM^out^, reporting stiffer asymmetric bilayers as compared to their scrambled systems (17). However, the stiffening effect was mainly dominating at lower temperatures (30 °C), i.e. within the chain melting regime, and decreased upon increasing temperature to 45°C, the highest studied temperature by these authors. It thus appears that the stiffening effect is abolished upon further increasing temperature, i.e. when reaching the here used 50 °C. The other two aLUVs with POPE enriched inner leaflets behaved much different. When either POPC or POPC/MSM mixtures were present in the outer leaflet, aLUVs were much stiffer than all other symmetric control LUVs, i.e. exceeding even the upper boundaries of inner/outer leaflet LUVs. The stiffening effect was slightly more expressed for POPE^in^/POPC^out^, with Δ*κ*_*r*_ ≡ *κ*^aLUV^/*κ*^scrambled^ ∼ 1.5, whereas Δ*κ*_*r*_ ∼ 1.4 for POPE^in^/(POPC/MSM)^out^.

Asymmetric membranes with POPE/POPS mixtures showed even larger rigidity differences between aLUVs and scrambled vesicles. The largest relative bending changes between these two systems occurred again in case of POPC outer leaflets with Δ*κ*_*r*_ ∼ 2.2, followed by (POPC/MSM)^out^ (Δ*κ*_*r*_ ∼ 1.9), and MSM^out^ (Δ*κ*_*r*_ ∼ 1.7). Further, the MSM^out^ aLUVs had an about equal bending rigidity than symmetric MSM-enriched outer leaflet vesicles. Asymmetric membranes with either POPC or POPC/MSM outer leaflets in turn revealed bending rigidities much outside the *κ* boundaries set by the symmetric cognate leaflet vesicles.

## DISCUSSION

We observed anomalous stiffening for most of the presently studied asymmetric lipid membranes. Here anomalous refers to bilayers, which are even stiffer than the upper rigidity boundary set by their symmetric cognate leaflet vesicles. A similar effect was previously reported for asymmetric giant unilamellar vesicles composed of POPC and DOPC (14), as well as DOPC and dimyristoylphosphatidylcholine (16). In turn, Rickeard et al. (17) found that the *κ* values of POPE/ESM aLUVs matched those of outer leaflet symmetric vesicles for fluid bilayers. Thus, the bending rigidity of these asymmetric membranes was not anomalously stiffer than the symmetric controls, but dominated by the more rigid leaflets. We found a complete loss of this mechanical coupling for the same asymmetric membranes at slightly higher temperatures (Fig. 4), most likely due to interactions of entropic origin. Similarly, POPE/MSM aLUVs showed no anomalous coupling of its leaflets. In stark contrast, POPE/POPC asymmetric bilayers were about 1.5 times more rigid than the cognate leaflet systems, clearly signifying anomalous membrane stiffening. This effect was enhanced for aLUVs with POPE/POPS enriched inner leaflets, except for (POPE/POPS)^in^/MSM^out^, whose bending rigidity was dominated by the more rigid leaflet, following the above discussed behavior of POPE/ESM asymmetric membranes at lower temperatures (17).

Attempting to reconcile our experimental observations within existing theoretical frameworks we first checked whether the differences could be simply understood in terms of the polymer brush model (21), i.e. differences in membrane thickness. Indeed, we observed a thickening of aLUVs for most presently systems, which would agree with a decreased bending flexibility (Tab. 1). Assuming that the area expansion modulus is not affected by membrane asymmetry, Δ*κ*_*r*_, determined above from NSE experiments, can also be estimated using the *D*_HH_-values reported in Tabs. S4, S6, and S7. This yields Δ*κ*_*r*_ values between 0.9 and 1.2 for the presently studied samples, but with no correlation to the Δ*κ*_*r*_ values found from NSE data (Fig. 4). This indicates that the area expansion modulus of asymmetric membranes differs significantly from that of scrambled vesicles, which would agree with a previous report (16).

Hossein and Deserno (8, 13) recently predicted a stiffening transition in asymmetric membranes based on differential stress (nonvanishing lateral tension) between the two leaflets. Such differential stress can arise either from compositional asymmetry, or leaflet under/overcrowding (number asymmetry). The latter type of asymmetry is difficult to control experimentally. Experimental studies using cyclodextrin-mediated lipid exchange, including this one, typically report some amounts of donor lipid within the inner leaflet (7, 12, 17, 36, 41). This suggests a rapid equilibration of any differential stress resulting from number asymmetry. We note that anomalous stiffening for asymmetric bilayers was also reported using a very different sample preparation technique (14, 16). Anomalous membrane stiffening thus is unlikely a manufacturing artifact.On quantitative grounds, we can use the area per lipid determined by SANS/SAXS as indicators of leaflet under/overcrowding. Table 1 reports the relative area per lipid changes between asymmetric and scrambled vesicles. In case of significant stress due to lateral area mismatch between leaflets in asymmetric membranes, we would expect also to observe corresponding area per lipid changes upon lipid scrambling due to stress relaxation. Variations of Δ*A*/*A* are, however, within a few percent, and do not correlate with the observed effects on bending rigidity. Hence, we can rule out membrane stiffening arising from number asymmetry.

Focusing next on compositional asymmetry we estimate the differential stress in flat membranes (8)

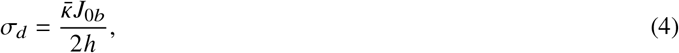

where *J*_0,*b*_ is the rigidity-weighted difference of the outer leaflet 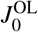 and inner leaflet 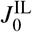 spontaneous curvatures 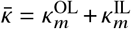 and is the leaflet averaged bending rigidity. Table 3 shows the results using previously reported spontaneous monolayer data (42, 43). The calculated *J*_0,*b*_’s are all close to 0 and have with the exception of (POPE/POPS)^in^/(POPC/MSM)^out^ all positive signs. Consequently, also differential stresses are rather small with absolute values varying between 0.16 mN/m and 0.95 mN/m. Correcting our *κ* for contributions of the tilt modulus (see above) would increase the reported *σ*_*d*_ values, but by not more than a factor of three, i.e. still yielding rather small *σ*_*d*_’s. Thus, contributions of compositional asymmetry to differential stress remain negligible, mostly due to the small *J*_0,*b*_ values. Moreover, relative changes of *σ*_*d*_ also do not explain the variation of *κ* for different samples and anomalous stiffening (Fig. 4). In particular aLUVs with the highest *κ* are among those with lowest |*σ*_*d*_ |.

**Table 3:** Elastic leaflet properties and resulting theoretical rigidity-weighted spontaneous curvature and differential stress for aLUVs.

| | $J_0^{\text{IL}}$ [nm<sup>-1</sup>]<sup>a</sup> | $J_0^{\text{OL}}$ [nm<sup>-1</sup>]<sup>a</sup> | $\kappa_m^{\text{IL}}$ [k<sub>B</sub>T]<sup>b</sup> | $\kappa_m^{\text{OL}}$ [k<sub>B</sub>T]<sup>b</sup> | $h$ [Å]<sup>c</sup> | $J_{0b}$ [nm<sup>-1</sup>] | $\sigma_d$ [mN/m] |
| --- | --- | --- | --- | --- | --- | --- | --- |
| POPE<sup>in</sup>/ESM<sup>out</sup> | -0.28 | -0.06 | 4.2 | 7.5 | 16.9 | 0.06 | 0.96 |
| POPE<sup>in</sup>/MSM<sup>out</sup> | -0.23 | -0.14 | 5.3 | 5.3 | 16.8 | 0.05 | 0.63 |
| POPE<sup>in</sup>/POPC<sup>out</sup> | -0.25 | -0.17 | 4.4 | 4.5 | 16.0 | 0.04 | 0.47 |
| POPE<sup>in</sup>/(POPC/MSM)<sup>out</sup> | -0.17 | -0.09 | 5 | 5.5 | 16.3 | 0.03 | 0.49 |
| (POPE/POPS)<sup>in</sup>/MSM<sup>out</sup> | -0.21 | -0.08 | 3.9 | 9.2 | 16.6 | 0.01 | 0.11 |
| (POPE/POPS)<sup>in</sup>/POPC<sup>out</sup> | -0.21 | -0.15 | 4.2 | 4.2 | 15.2 | 0.03 | 0.37 |
| (POPE/POPS)<sup>in</sup>/(POPC/MSM)<sup>out</sup> | -0.14 | -0.15 | 4.5 | 5 | 16.1 | -0.01 | -0.17 |

<sup>a</sup> Data taken from ref. (42, 43) and averaged according to molecular leaflet composition. Intrinsic curvatures of MSM, ESM and POPS were assumed to be zero.
<sup>b</sup> Calculated from symmetric reference LUVs for inner and outer leaflets using $\kappa_m = \kappa/2$ .
<sup>c</sup> Average value for both leaflets.

For the sake of the argument — and in absence of any theory we are aware of — let us therefore speculate on the putative role of fluctuations. The undulatory motions of the fluctuation spectrum of lipid membranes comprise of bending and protrusion modes (44, 45). Here, we probed bending modes, which dominate the fluctuation spectrum for distances larger than the membrane thickness. At shorter distances these interleaflet dynamics are thought be essentially decoupled. In this ‘protrusion regime’ shorter wavelengths fluctuations (alternatively hard modes) are dominated by single lipid motions (e.g. protrusions). While this theory agrees well with large scale molecular dynamics simulations of symmetric bilayers (44, 45) it is less clear if it applies also to asymmetric membranes. For instance, our data might suggest that fluctuations in some of our aLUVs are coupled even at distances smaller than the membrane thickness, increasing the weight of ‘hard’ modes on bending fluctuations. This enhanced intra leaflet coupling appears to particularly occur for POPC-enriched outer leaflets. Both, POPE and POPS carry primary amines in their headgroups, enabling intermolecular H-bond formation, which is indeed known to stabilize bilayer structure (46). The headgroup charge of POPS additionally seems to accentuate the intraleaflet coupling, possibly by giving rise to a transmembrane potential, or by increasing contributions from surface tension. It would be interesting to modulate in future experiments their contribution by changing the pH or ionic strength of the aqueous phase.

Interestingly, no anomalous elastic coupling was observed for ESM and MSM-enriched outer leaflets and the stiffening effect was partially reduced for POPC/MSM mixtures. ESM and MSM are natural lipid extracts, containing also some long acyl chains. In particular, MSM is highly chain-asymmetric with mostly 22:0, 23:0 and 24:0 hydrocarbons and thus prone to interdigitation (12). The hydrocarbon chain composition of ESM is dominated by 16:0 hydrocarbons, but also contains few longer hydrocarbons which can interdigitate into the opposing leaflet. Hydrocarbon interdigitation of POPC in turn is negligible (24). Consequently, our data suggest that the anomalous mechanical coupling induced by the enrichment of aminophospholipids in one leaflet can be alleviated by hydrocarbon interdigitation.

The potential biological relevance is as intriguing as our findings. Several cellular processes, occurring either at large scales, such as the formation of exo-, or endosomes, or within the single molecular regime, e.g. transport or signalling events, depend on membrane elasticity (4, 5). By tuning the leaflet distribution of lipids in given regions of the plasma membrane through an orchestrated operation of flipases/flopases and scramblases cells might control its elasticity over a broad range of bending rigidites and exploit its effect on certain cellular processes. Indeed, there appears to be experimental evidence for softer scrambled plasma membranes derived from live cell as compared to their native states (Ilya Levental, personal communication). Further scrutiny of the here reported systems, including cholesterol and different buffer conditions is challenging but certainly warranted.

## CONCLUSION

We observed that asymmetric transbilayer distribution of the aminophospholipids POPE and POPS can lead to an anomalous stiffing of membranes, i.e. exceeding not only the bending rigidity of their scrambled analogs, but also that of their cognate leaflets in symmetric bilayers. These effects can be alleviated at least in part by the addition of sphingomyelin in the outer leaflet, possibly due to hydrocarbon chain interdigitation. We speculate that an asymmetric distribution of lipids with specific properties (charge, H-bonding abilities) leads to an enhanced mechanical coupling of membranes, by increasing the weight of hard undulatory modes through intraleaflet interactions. This strongly encourages theoretical and computational studies along those directions. Although the presently studied systems still lack important components of mammalian plasma membranes (e.g. cholesterol) it is plausible that cells have evolved to a state where they can exploit a large range of bending rigidities for protein function by dynamically modulating leaflet-specific lipid composition.

## AUTHOR CONTRIBUTIONS

MPKF, YG, LP and GP designed the research. MPKF, PP, EFS and OC carried out all experiments. MPKF analyzed the data. MPKF and GP wrote the article.

## ACKNOWLEDGMENTS

This research was supported by the ILL graduate school, Ph.D. No. 181-19. PP and EFS are supported by the Austrian Science Fund (FWF), grant no. P32514 to GP. We thank the ILL and the ESRF for beamtime, the PSCM for access to support laboratories, and Petra Pernot, Mark Tully, Dihia Moussaoui and Anton Popov for their assistance with experiments at BM29 and Krishna Batchu for help with GC measurements. We thank E. London, H. Heerklotz, F. Heberle, M. Doktorova and M. Deserno for constructive discussions.

## SUPPLEMENTARY MATERIAL

An online supplement to this article can be found by visiting BJ Online at http://www.biophysj.org.

