## Supplementary Information for "Distributing Aminophospholipids Asymmetrically Across Leaflets Causes Anomalous Membrane Stiffening"

#### Contents

**1 Evaluation of NMR data** **2**

**2 SDP-model used for SAS-analysis** **3**

**3 The contribution of diffusion in NSE-data modelling** **6**

**4 Bending Rigidities of Lipid mixtures** **7**

**5 SAS-model parameters of donor/acceptor lipids** **8**

**6 SAS-model parameters of asymmetric vesicles** **10**

**7 SAS-model parameters of reference LUVs** **15**

### 1 Evaluation of NMR data

Figure S1 shows an exemplary NMR spectrum of asymmetric vesicles before and after addition of $Pr^{3+}$. The areas $A_0$ and $A_1$ under the Lorentzian fits of the peak at 3.4 ppm (dotted lines) correspond to the total amount of choline headgroups (before) and the amount of choline lipids inside the vesicle (after), respectively. The mol-fraction of PC-lipids (POPC and/or Sphingomyelin) in the inner leaflet is therefore given by $\chi_P^{in} = A_1/A_0$.

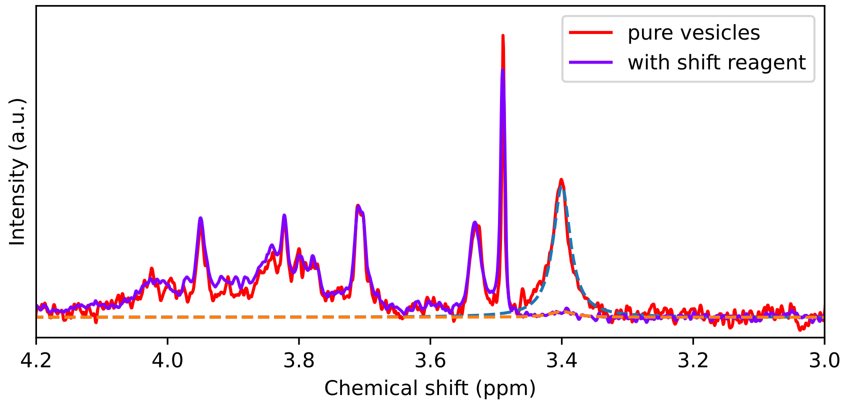


Figure S1: The graph shows a part of the NMR spectrum of asymmetric vesicles before and after adding $Pr^{3+}$. While the majority of the choline peak is moved from its original position (3.4 ppm), the rest of the spectrum is unaffected. Dotted lines mark Lorentzians used to fit the peaks and to calculate the underlying areas.

#### 2 SDP-model used for SAS-analysis

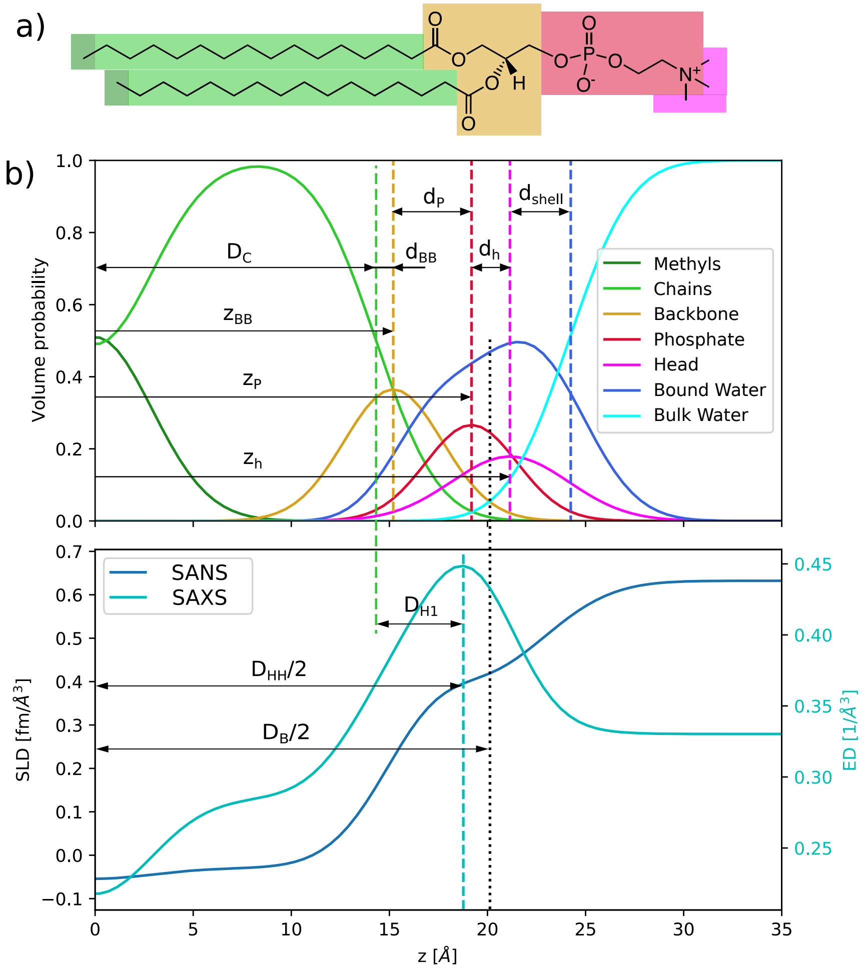


Figure S2: Lipid parsing and scattering length density profile modelling: a) The upper graphic shows how a lipid is divided in fragments on the example of DPPC. The parsing off all other used lipids can be found in Table S1. b) The color-coded arrangement of functions (*pdf*) in the upper diagram corresponds to the probability density to find a fragment at a distance $z$ from the bilayer center. The *pdf*s for the terminal methyls ($T$), lipid backbones ($BB$), phosphate groups ($P$) and heads ($h$) are modelled by Gaussian functions. Hydrocarbon chains ($HC$) and a hydration water layer (bound water, $BW$) are slabs with altitude 1 and smeared with error-functions. From these, the functions of $T$ and $BB + P + H$ are substracted respectively. The *pdf* of the solvent (bulk water $W$) fills the remaining space, so that the sum of all *pdf*s is 1 at all $z$. The lower diagram shows the neutron scattering length density ($SLD$) and electron density ($ED$) profiles of the modelled lipid in $D_2O$. It is obtained from the sum over all *pdf*s, multiplied with the respective $SLD$ or $ED$.

Table S1: List of lipid fragments and chemical compositions for all used lipids. X denote exchangable hydrogens, which are replaced with deuterium in a D2O-environment.

| Fragment | Terminal methyls | Chains without T | Backbone | Phosphate group | Head |
|---|---|---|---|---|---|
| Abbr. | T | HC | BB | P | h |
| DPPC | $(CH_3)_2$ | $(CH_2)_{28}$ | $C_5O_4H_5$ | $PO_4(CH_2)_2N$ | $(CH_3)_3$ |
| DPPG | $(CH_3)_2$ | $(CH_2)_{28}$ | $C_5O_4H_5$ | $PO_4$ | $C_3H_5O_2X_2$ |
| POPE | $(CH_3)_2$ | $(CH_2)_{28}(CH)_2$ | $C_5O_4H_5$ | $PO_4$ | $(CH_2)_2NX_3$ |
| POPG | $(CH_3)_2$ | $(CH_2)_{28}(CH)_2$ | $C_5O_4H_5$ | $PO_4$ | $C_3H_5O_2X_2$ |
| POPS | $(CH_3)_2$ | $(CH_2)_{28}(CH)_2$ | $C_5O_4H_5$ | $PO_4$ | $C_3H_2X_3NO_2H$ |
| POPC | $(CH_3)_2$ | $(CH_2)_{28}(CH)_2$ | $C_5O_4H_5$ | $PO_4(CH_2)_2N$ | $(CH_3)_3$ |
| ESM | $(CH_3)_2$ | $(CH_2)_{26.6}(CH)_2$ | $C_4O_2NH_6$ | $PO_4(CH_2)_2N$ | $(CH_3)_3$ |
| MSM | $(CH_3)_2$ | $(CH_2)_{32}(CH)_2$ | $C_4O_2NH_6$ | $PO_4(CH_2)_2N$ | $(CH_3)_3$ |

SAS-data are modelled using the vesicle form factor $F_{sphere}$, the bilayer form $F_{bil}$, which we split into real and imaginary part, and the incoherent background $I_{inc}$. Furthermore, we applied a Gaussian polydispersity on the hydrophobic thickness $D_C$, which results in a weighted average over several bilayer form factors $F_{bil,k}$. A more detailed description of the model was published earlier [1]. Due to the lack of contrast between both leaflets, we are not able to locate the bilayer center by SANS and therefore fixed the position of the terminal methyl group $z_T = 0$ in this study. For all symmetric references, the imaginary part of the bilayer form factor is equal 0 and only one leaflet has to be modelled. All quantities $\Delta\rho$ denote the scattering length density contrast of moiety $k$ with respect to the water/heavy water environment $\Delta\rho_k = \rho_k - \rho_W$. Distances are defined in Fig. S2, where $D_B$ is defined as the Luzzati bilayer thickness and $D_{HH}$ the distance between the maxima in the ED-profile. $D_{H1} = (D_B - D_{HH})/2$. $\sigma_k$ are the standard deviations of the respective *pdf*s. $d_{shell}$ was fixed to 3.1 Å for all samples. Further parameters: $A$ area per lipid; $V_L$ total lipid volume; $V_H$ total head group volume (back bone, phosphate group and head); $r_{BB} = V_{BB}/V_H$, $r_P = V_P/V_H$, $r = V_{CH3}/V_{CH2}$, $r_{12} = V_{CH}/V_{CH2}$; $V_{W,bound}$ volume per bound water molecule; $n_W$ number of bound water molecules; $Y$ relative interdigitation (see [2]).

$$I(q) \propto F_{sphere}(R_m, \sigma_R) \sum_k \mathcal{N}(D_{C,k}|\bar{D_C}, \sigma_{\rm poly}) \left\{ |F_{bil,k}^{real}|^2 + |F_{bil,k}^{imag}|^2 | \right\} + I_{inc} \tag{1}$$

with

$$\begin{aligned} F_{bil,k}^{real} =& 4\Delta\rho_{\rm bw}\frac{1}{q}e^{-\frac{q^2\sigma_{\rm CH2}^{\rm in\,2}}{2}} \sin\left(q\frac{d_{\rm BB}^{\rm in} + d_P^{\rm in} + d_{shell}^{\rm in}}{2}\right) \\ & \cos\left(q(-D_{C,k}^{\rm in} - \frac{d_{\rm BB}^{\rm in} + d_P^{\rm in} d_{shell}^{\rm in}}{2})\right) + \\ & 2(\Delta\rho_P^{\rm in} - \Delta\rho_{\rm bw})\frac{V_P^{in}}{A}e^{-\frac{q^2\sigma_P^{\rm in\,2}}{2}} \cos\left(q(-D_{C,k}^{\rm in} - d_{\rm BB}^{\rm in} - d_P^{\rm in}/2)\right) + \\ & 2(\Delta\rho_{\rm BB}^{\rm in} - \Delta\rho_{\rm bw})\frac{V_{\rm BB}^{\rm in}}{A}e^{-\frac{q^2\sigma_{\rm BB}^{\rm in\,2}}{2}} \cos\left(q(-D_{C,k}^{\rm in} - d_{\rm BB}^{\rm in}/2)\right) + \\ & 2\Delta\rho_{\rm CH2}^{\rm in}\frac{1}{q}e^{-\frac{q^2\sigma_{\rm CH2}^{\rm in\,2}}{2}} \sin\left(qD_{C,k}^{\rm in}/2\right) \cos\left(q(-D_{C,k}^{\rm in}/2\right) + \\ & 2\Delta\rho_{\rm CH2}\frac{1}{q}e^{-\frac{q^2\sigma_{\rm CH2}^2}{2}} \sin\left(qD_{C,k}^{\rm out}/2\right) \cos\left(qD_{C,k}^{\rm out}/2\right) + \\ & 2(\Delta\rho_{\rm BB}^{\rm out} - \Delta\rho_{\rm bw})\frac{V_{\rm BB}^{\rm out}}{A}e^{-\frac{q^2\sigma_{\rm BB}^{\rm out\,2}}{2}} \cos\left(q(D_{C,k}^{\rm out} + d_{\rm BB}^{\rm out}/2)\right) + \\ & 2(\Delta\rho_P^{\rm out} - \Delta\rho_{\rm bw})\frac{V_P^{\rm out}}{A}e^{-\frac{q^2\sigma_P^{\rm out\,2}}{2}} \cos\left(q(D_{C,k}^{\rm out} + d_{\rm BB}^{\rm out} + d_P^{\rm out}/2)\right) + \\ & 4\Delta\rho_{\rm bw}\frac{1}{q}e^{-\frac{q^2\sigma_{\rm CH2}^{\rm out\,2}}{2}} \sin\left(q\frac{d_{\rm BB}^{\rm out} + d_P^{\rm out} + d_{shell}^{\rm out}}{2}\right) \\ & \cos\left(q(D_{C,k}^{\rm out} + \frac{d_{\rm BB}^{\rm out} + d_P^{\rm out} + d_{shell}^{\rm out}}{2})\right) \end{aligned} \tag{2}$$

and

$$
\begin{aligned}
F^{imag}_{bil,k} =& 4\Delta\rho_{\mathrm{bw}}\frac{1}{q}\mathrm{e}^{-\frac{q^2\sigma^{\mathrm{in}\,2}_{\mathrm{CH2}}}{2}}\sin\left(q\frac{d^{\mathrm{in}}_{\mathrm{BB}}+d^{\mathrm{in}}_{P}+d^{\mathrm{in}}_{shell}}{2}\right) \\
& \sin\left(q(-D^{\mathrm{in}}_{C,k}-\frac{d^{\mathrm{in}}_{\mathrm{BB}}+d^{\mathrm{in}}_{P}d^{\mathrm{in}}_{shell}}{2})\right)+ \\
& 2(\Delta\rho^{\mathrm{in}}_{P}-\Delta\rho_{\mathrm{bw}})\frac{V^{in}_{P}}{A}\mathrm{e}^{-\frac{q^2\sigma^{\mathrm{in}\,2}_{P}}{2}}\sin\left(q(-D^{\mathrm{in}}_{C,k}-d^{\mathrm{in}}_{\mathrm{BB}}-d^{\mathrm{in}}_{P}/2)\right)+ \\
& 2(\Delta\rho^{\mathrm{in}}_{\mathrm{BB}}-\Delta\rho_{\mathrm{bw}})\frac{V^{\mathrm{in}}_{\mathrm{BB}}}{A}\mathrm{e}^{-\frac{q^2\sigma^{\mathrm{in}\,2}_{\mathrm{BB}}}{2}}\sin\left(q(-D^{\mathrm{in}}_{C,k}-d^{\mathrm{in}}_{\mathrm{BB}}/2)\right)+ \\
& 2\Delta\rho^{\mathrm{in}}_{\mathrm{CH2}}\frac{1}{q}\mathrm{e}^{-\frac{q^2\sigma^{\mathrm{in}\,2}_{\mathrm{CH2}}}{2}}\sin\left(qD^{\mathrm{in}}_{C,k}/2\right)\sin\left(q(-D^{\mathrm{in}}_{C,k}/2\right)+ \\
& \frac{\Delta\rho^{\mathrm{in}}_{\mathrm{T}}+\Delta\rho^{\mathrm{out}}_{\mathrm{T}}}{2}\mathrm{e}^{-\frac{q^2\sigma^2_{\mathrm{T}}}{2}}+ \\
& 2\Delta\rho_{\mathrm{CH2}}\frac{1}{q}\mathrm{e}^{-\frac{q^2\sigma^2_{\mathrm{CH2}}}{2}}\sin\left(qD^{\mathrm{out}}_{C,k}/2\right)\sin\left(qD^{\mathrm{out}}_{C,k}/2\right)+ \\
& 2(\Delta\rho^{\mathrm{out}}_{\mathrm{BB}}-\Delta\rho_{\mathrm{bw}})\frac{V^{\mathrm{out}}_{\mathrm{BB}}}{A}\mathrm{e}^{-\frac{q^2\sigma^{\mathrm{out}\,2}_{\mathrm{BB}}}{2}}\sin\left(q(D^{\mathrm{out}}_{C,k}+d^{\mathrm{out}}_{\mathrm{BB}}/2)\right)+ \\
& 2(\Delta\rho^{\mathrm{out}}_{P}-\Delta\rho_{\mathrm{bw}})\frac{V^{\mathrm{out}}_{P}}{A}\mathrm{e}^{-\frac{q^2\sigma^{\mathrm{out}\,2}_{P}}{2}}\sin\left(q(D^{\mathrm{out}}_{C,k}+d^{\mathrm{out}}_{\mathrm{BB}}+d^{\mathrm{out}}_{P}/2)\right)+ \\
& 4\Delta\rho_{\mathrm{bw}}\frac{1}{q}\mathrm{e}^{-\frac{q^2\sigma^{\mathrm{out}\,2}_{\mathrm{CH2}}}{2}}\sin\left(q\frac{d^{\mathrm{out}}_{\mathrm{BB}}+d^{\mathrm{out}}_{P}+d^{\mathrm{out}}_{shell}}{2}\right) \\
& \sin\left(q(D^{\mathrm{out}}_{C,k}+\frac{d^{\mathrm{out}}_{\mathrm{BB}}+d^{\mathrm{out}}_{P}+d^{\mathrm{out}}_{shell}}{2})\right)
\end{aligned} \tag{3}
$$

To describe the contribution from the overall vesicle shape we use the Schultz-distributed form factor of a sphere, as described in [3]:

$$
F_{sphere} = \frac{8\pi^2(z+1)(z+2)}{s^2q^2}\left\{1-\left(1+\frac{4q^2}{s^2}\right)^{-(z+3)/2}\cos\left[(z+3)\arctan\left(\frac{2q}{s}\right)\right]\right\} \tag{4}
$$

Mean vesicle radius $R_m$ and polydispersity $\sigma_R$ enter via the auxiliary quantities $s = \frac{R_m}{\sigma_R^2}$ and $z = \frac{R_m^2}{\sigma_R^2} - 1$.

#### 3 The contribution of diffusion in NSE-data modelling

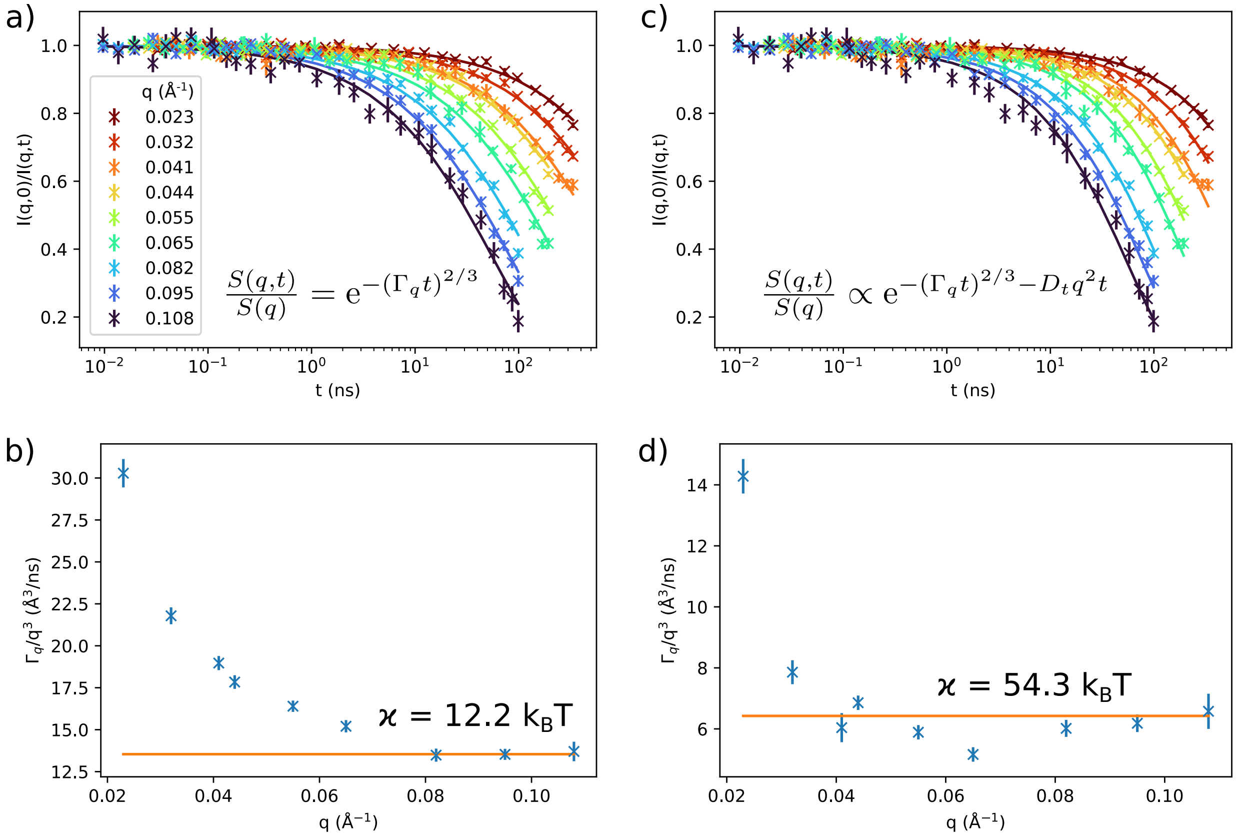


Figure S3: The graphic shows the comparison of 2 models used to evaluate NSE-data of DPPC-vesicles at 50 °C: (I) the pure Zilman-Granek (ZG) model (a,b) and (II) the ZG model with a contribution from diffusion (c,d), using a translational diffusion constant $D_t = 0.66$ Å$^2$/ns, measured by dynamic light scattering. The formulas to fit data are given in (a) and (c). Both models are in reasonable agreement with the raw data, however using model (I), the resulting values for the $q$-dependent decay constant $\Gamma_q$ (b) do not follow $q^3$ as predicted by the ZG-theory. Model (II) improves the agreement over a wider $q$-range (d), but still deviates at low $q$. The given values for the bending rigidity $\kappa$ in (b) and (d) correspond to the orange lines.

### 4 Bending Rigidities of Lipid mixtures

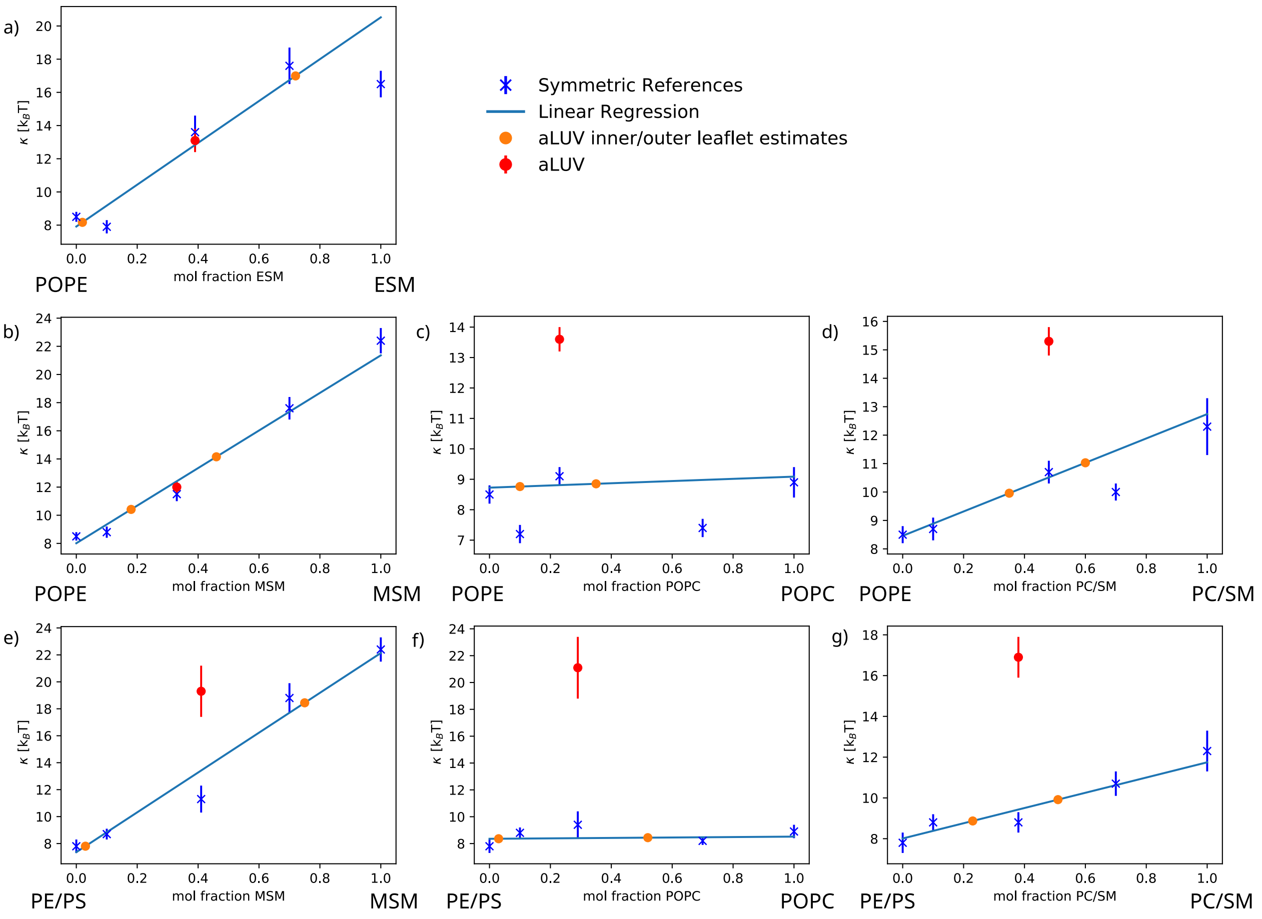


Figure S4: The graphic shows the measured bending rigidities $\kappa$ for all aLUVs (red) as well as symmetric references (blue). We performed linear regressions through the reference samples and extrapolated to retrieve the $\kappa$ at the measured inner/outer leaflet compositions (orange). Some points had to be excluded from the analysis due to large deviations from the linear law: a) $X_{MSM} = 0.46$, b) $X_{ESM} = 0.1$ and $X_{ESM} = 1$, d) $X_{POPC} = 0.1$ and $X_{POPC} = 0.7$, e) $X_{PC/SM} = 0.7$. The origin of these deviations is unclear. In the case of a) the difference of both asymmetric and symmetric sample suggests that the donor-acceptor exchange was lower than expected from the previously prepared similar sample [1].

### 5 SAS-model parameters of donor/acceptor lipids

Table S2: Summary of structural properties of symmetric LUVs at 50°C.

| | A (Å$^2$) | $D_{\mathrm{HH}}$ (Å)[c] | $2D_C$ (Å) | $h$ (Å) |
|---|---|---|---|---|
| DPPC[a] | 63.1 ± 1.3 | 37.5 ± 1.1 | 28.6 ± 0.9 | 15.2 ± 1.2 |
| POPC[a] | 67.5 ± 1.4 | 37.5 ± 1.1 | 28.4 ± 0.9 | 14.8 ± 1.2 |
| MSM[a] | 64.8 ± 1.3 | 43.0 ± 1.1 | 32.8 ± 1.0 | 18.4 ± 1.5 |
| ESM | 56.2 ± 1.1 | 41.5 ± 1.1 | 33.6 ± 1.0 | 19.8 ± 1.6 |
| POPE | 60.5 ± 1.2 | 36.0 ± 1.1 | 31.2 ± 0.9 | 17.0 ± 1.4 |
| POPE/POPS[b] | 65.3 ± 1.3 | 37.1 ± 1.1 | 28.9 ± 0.9 | 17.0 ± 1.4 |

[a] Structural data taken from ref. [2].
[b] 7:3 mol/mol.
[c] head-to-headgroup distance.

Table S3: Properties of symmetric lipid bilayers from SAXS/SANS analysis at 50 °C containing POPE/POPG 9:1 (POPE), POPE/POPS 7:3 (PE/PS) and ESM/DPPG 19:1 (ESM).

| | $\epsilon$ [%] | POPE | PE/PS | ESM |
|---|---|---|---|---|
| $V_L^*$ [Å$^3$] | | 1193.8 | 1199.3 | 1218.5 |
| $V_H^*$ [Å$^3$] | | 249.6 | 254.9 | 274.9 |
| $r_{BB}^*$ | | 0.51 | 0.50 | 0.33 |
| $r_P^*$ | | 0.14 | 0.18 | 0.31 |
| $r^*$ | | 2.09 | 2.09 | 2.09 |
| $r_{12}^*$ | | 0.8 | 0.8 | 0.8 |
| $D_B$ [Å] | 3 | 39.5 | 36.8 | 43.4 |
| $D_{HH}$ [Å] | 3 | 36.0 | 37.1 | 41.5 |
| $2D_C$ [Å] | 3 | 31.2 | 28.9 | 33.6 |
| $D_{H1}$ [Å] | 20 | 2.4 | 4.1 | 4.0 |
| $A$ [Å$^2$] | 2 | 60.5 | 65.3 | 56.2 |
| $z_{BB}$ [Å] | 8 | 17.0 | 17.0 | 19.8 |
| $\sigma_{BB}$ [Å] | 20 | 2.5 | 2.5 | 3.2 |
| $z_P$ [Å] | 8 | 18.8 | 20.0 | 21.1 |
| $\sigma_P$ [Å] | 20 | 3.5 | 3.0 | 2.9 |
| $z_h$ [Å] | 3 | 23.8 | 20.0 | 21 |
| $\sigma_h^\dagger$ [Å] | | 2.8 | 3 | 3.0 |
| $\sigma_{HC}$ [Å] | | 2.5 | 2.5 | 2.5 |
| $\sigma_T$ [Å] | 5 | 3.0 | 3.0 | 4.1 |
| $\sigma_{poly}$ [%] | 6 | 4.0 | 8.1 | 3.5 |
| $V_{W,bound}$ [Å$^3$] | 6 | 29.6 | 29.4 | 27.3 |
| $n_W$ | 6 | 13.8 | 10.0 | 4.4 |
| $Y$ | 9 | 0.40 | 0.45 | 0.62 |

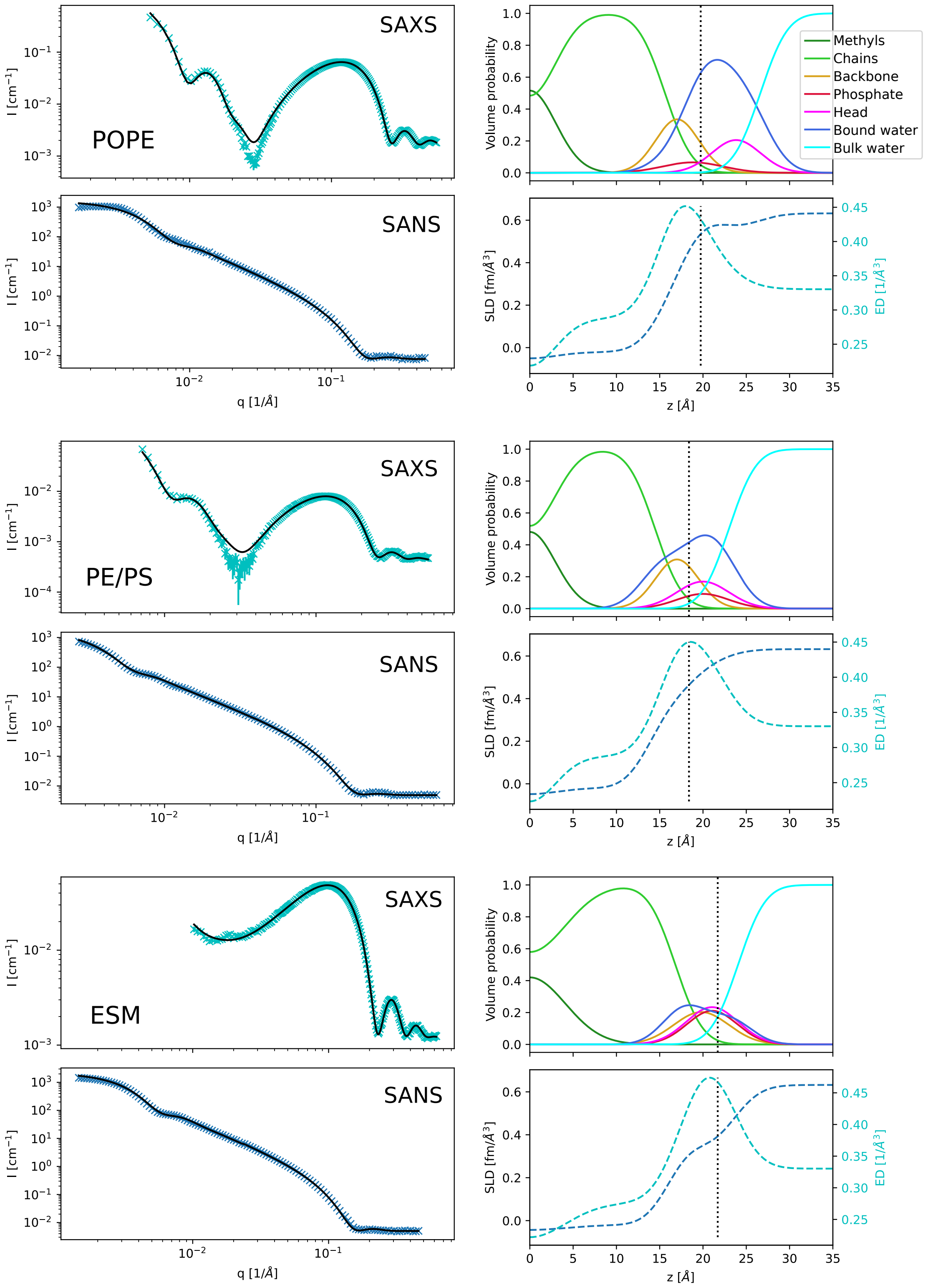


Figure S5: SAXS and SANS data with fits (black lines); SDP volume probability, electron density and neutron scattering length density profiles for symmetric lipid vesicles containing POPE/POPG 9:1, POPE/POPS 7:3 and Egg-SM/DPPG 19:1.

### 6 SAS-model parameters of asymmetric vesicles

Table S4: SAS-fitting parameters of aLUVs containing either a POPE/POPG 9:1 mixture (POPE) or a POPE/POPS 7:3 mixture (PE/PS) as acceptor lipids and ESM, POPC, MSM or a 1:1 POPC/MSM mixture (PC/SM) as donor lipids.

| | $\epsilon$ [%] | POPE ESM | POPE POPC | POPE MSM | POPE PC/SM | PE/PS POPC | PE/PS MSM | PE/PS PC/SM |
|---|---|---|---|---|---|---|---|---|
| Total acc/don % | 5 | 61:39 | 77:23 | 67:33 | 52:48 | 71:29 | 59:41 | 62:38 |
| In acc/don % | 5 | 98:2 | 90:10 | 82:18 | 65:35 | 97:3 | 97:3 | 77:23 |
| Out acc/don % | 5 | 28:72 | 65:35 | 54:46 | 40:60 | 48:52 | 25:75 | 49:51 |
| $D_B$ [Å] | 3 | 37.6 | 39.0 | 40.3 | 39.6 | 37.1 | 40.0 | 39.1 |
| $D_{HH}$ [Å] | 3 | 37.3 | 36.6 | 37.8 | 37.8 | 34.3 | 37.4 | 38.5 |
| $2D_C$ [Å] | 3 | 29.5 | 30.4 | 31.9 | 30.9 | 28.8 | 31.6 | 30.5 |
| $D_C^{in}$ [Å] | 5 | 14.3 | 15.2 | 15.4 | 15.7 | 13.5 | 16.3 | 15.1 |
| $D_C^{out}$ [Å] | 5 | 15.2 | 15.2 | 16.5 | 15.2 | 15.3 | 15.3 | 15.5 |
| $D_{H1}^{in}$ [Å] | 20 | 4.6 | 2.8 | 3.3 | 3.8 | 2.3 | 3.4 | 3.9 |
| $D_{H1}^{out}$ [Å] | 20 | 3.2 | 3.4 | 2.6 | 3.2 | 3.2 | 2.4 | 4.1 |
| $A^{av}$ [Å$^2$] | 2 | 64.0 | 62.0 | 61.4 | 62.9 | 66.0 | 62.8 | 63.4 |
| $z_{BB}^{in}$ [Å] | 6 | -16.4 | -16.0 | -16.2 | -16.5 | -14.3 | -17.1 | -15.9 |
| $z_{BB}^{out}$ [Å] | 6 | 17.4 | 16.0 | 17.3 | 16.0 | 16.1 | 16.1 | 16.3 |
| $\sigma_{BB}^{in/out}$ [Å] | | 2.5 | 2.5 | 2.5 | 2.5 | 2.5 | 2.5 | 2.5 |
| $z_P^{in}$ [Å] | 10 | -19.4 | -21.0 | -19.2 | -21.2 | -17.3 | -20.1 | -19.3 |
| $z_P^{out}$ [Å] | 10 | 22.4 | 19.0 | 21.5 | 19.0 | 19.7 | 19.9 | 20 |
| $\sigma_P^{in}$ [Å] | 10 | 2.0 | 3.5 | 2 | 3.0 | 4.0 | 2.0 | 2 |
| $\sigma_P^{out}$ [Å] | 10 | 4.0 | 2.0 | 3.5 | 2.3 | 3.0 | 3.8 | 2 |
| $z_h^{in}$ [Å] | 10 | -21.7 | -24.0 | -22.2 | -24.2 | -20.3 | -23.1 | -22.3 |
| $z_h^{out}$ [Å] | 10 | 24.6 | 22.0 | 24.5 | 22.0 | 22.7 | 22.9 | 23 |
| $\sigma_h^{in/out\dagger}$ [Å] | | 3.0 | 3.0 | 3 | 3.0 | 3.0 | 3.0 | 3 |
| $\sigma_{HC}^{in/out}$ [Å] | | 2.5 | 2.5 | 2.5 | 2.5 | 2.5 | 2.5 | 2.5 |
| $\sigma_T$ [Å] | 20 | 3.7 | 3.0 | 3.4 | 3.7 | 5.0 | 3.6 | 3.1 |
| $\sigma_{poly}$ [%] | 6 | 0.0 | 3.9 | 4.3 | 5.0 | 10.0 | 7.0 | 7 |
| $V_{W,bound}$ [Å$^3$] | 6 | 30.3 | 30.3 | 30.2 | 30.0 | 29.4 | 29.6 | 29.3 |
| $n_W^{in}$ | 6 | 14.4 | 15.4 | 11.8 | 14.6 | 14.2 | 10.3 | 12.8 |
| $n_W^{out}$ | 6 | 16.7 | 11.0 | 13.4 | 11.5 | 11.6 | 14.8 | 12.7 |
| $R_m$ [Å] | 10 | 361 | 361 | 345.3 | 373.4 | 369 | 361.2 | 381.9 |
| $\sigma_R$ [Å] | 10 | 133 | 133 | 101.4 | 111.7 | 116.1 | 133 | 134.7 |

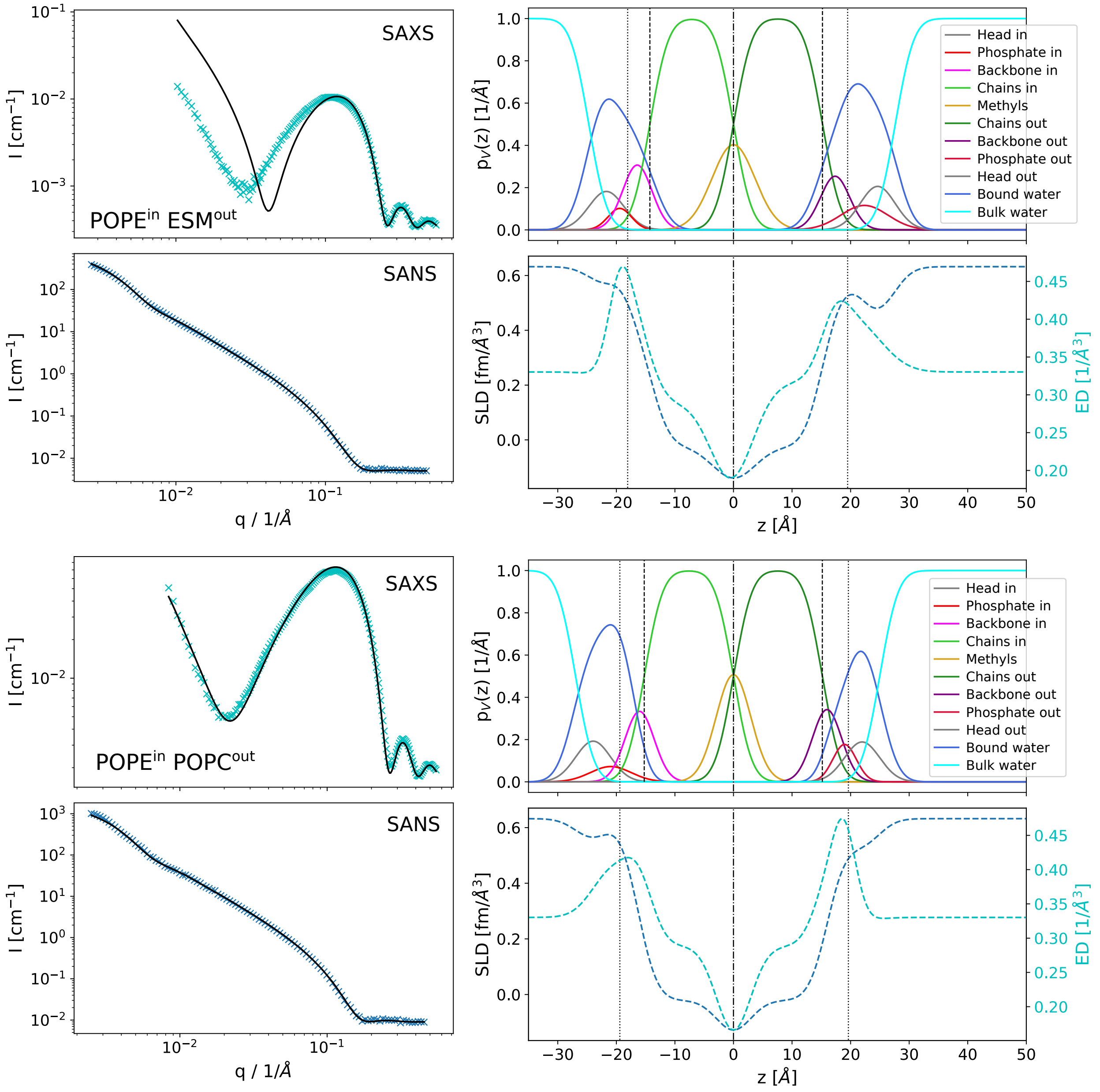


Figure S6: SAXS and SANS data with fits (black lines); SDP volume probability, electron density and neutron scattering length density profiles for the systems $POPE^{in}/ESM^{out}$ and $POPE^{in}/POPC^{out}$.

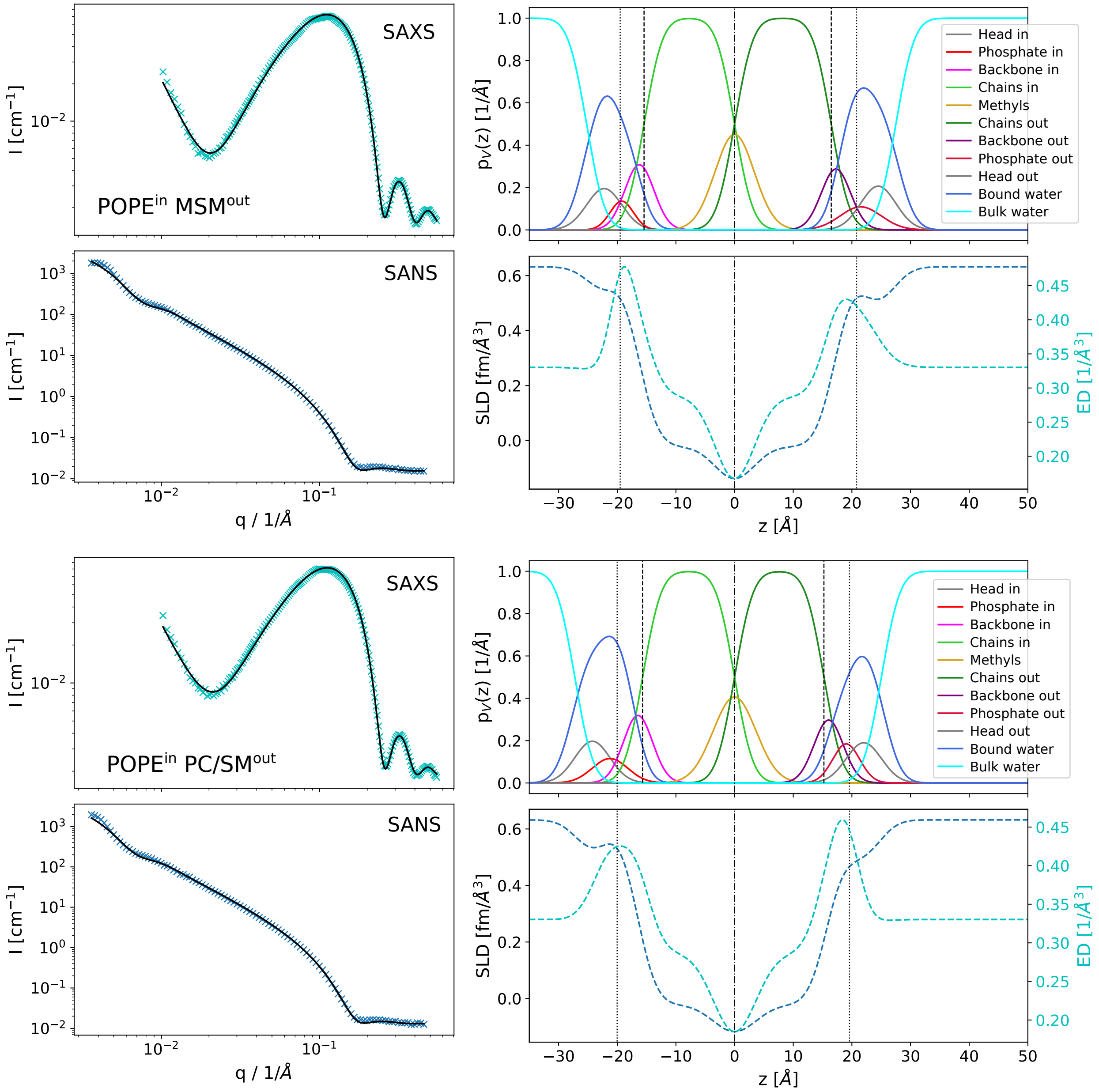


Figure S7: SAXS and SANS data with fits (black lines); SDP volume probability, electron density and neutron scattering length density profiles for the systems POPE$^{in}$/MSM$^{out}$ and POPE$^{in}$/(PC/SM)$^{out}$.

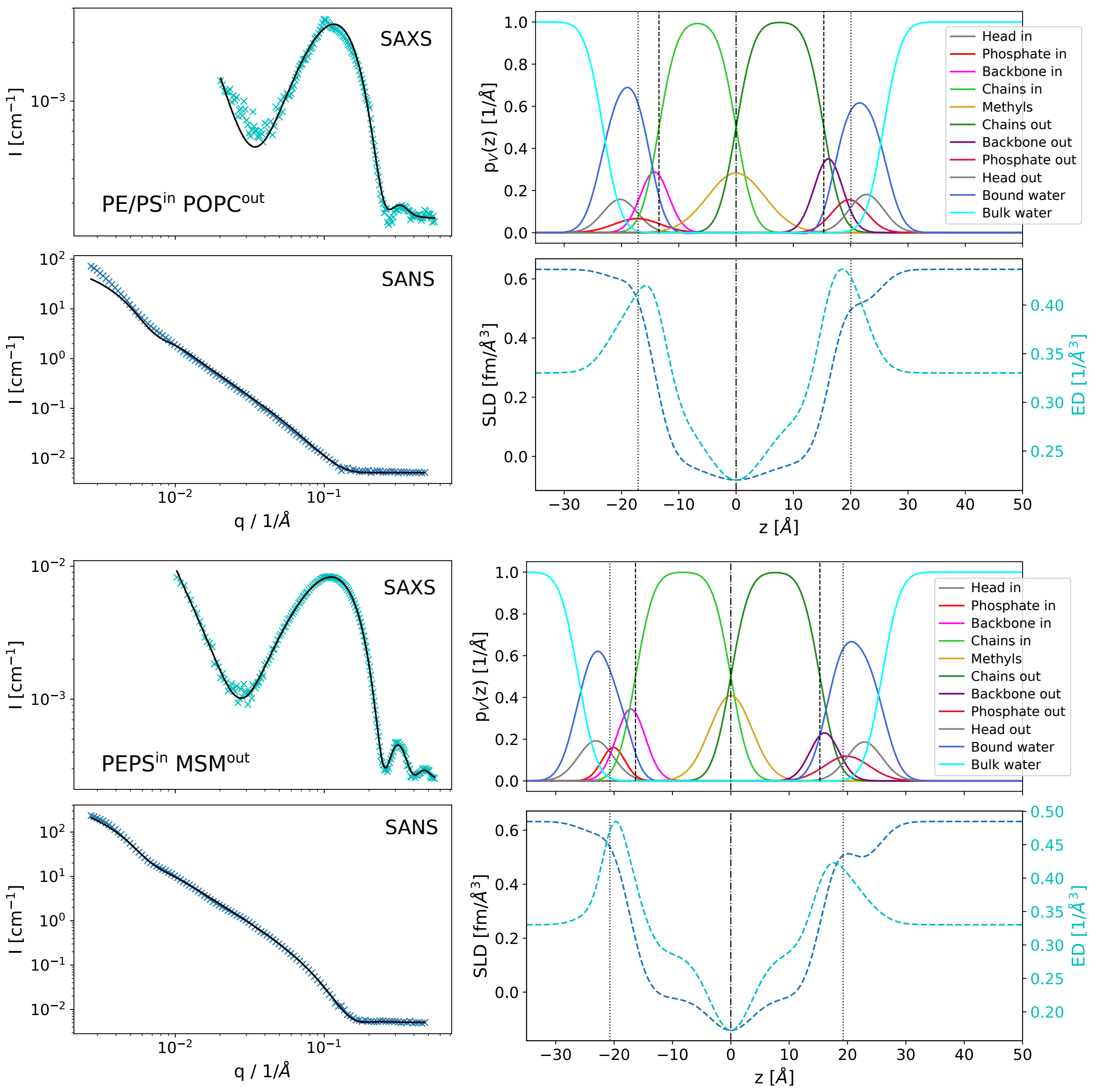


Figure S8: SAXS and SANS data with fits (black lines); SDP volume probability, electron density and neutron scattering length density profiles for the systems $(PE/PS)^{in}/POPC^{out}$ and $(PE/PS)^{in}/MSM^{out}$. SAXS data in the upper panel show a small bragg peak, which is probably connected to a small contamination of multilamellar donor vesicles in the sample. This could be also responsable for the disagreement between data and model in low-$q$ SANS. We expect a higher experimental error for this sample.

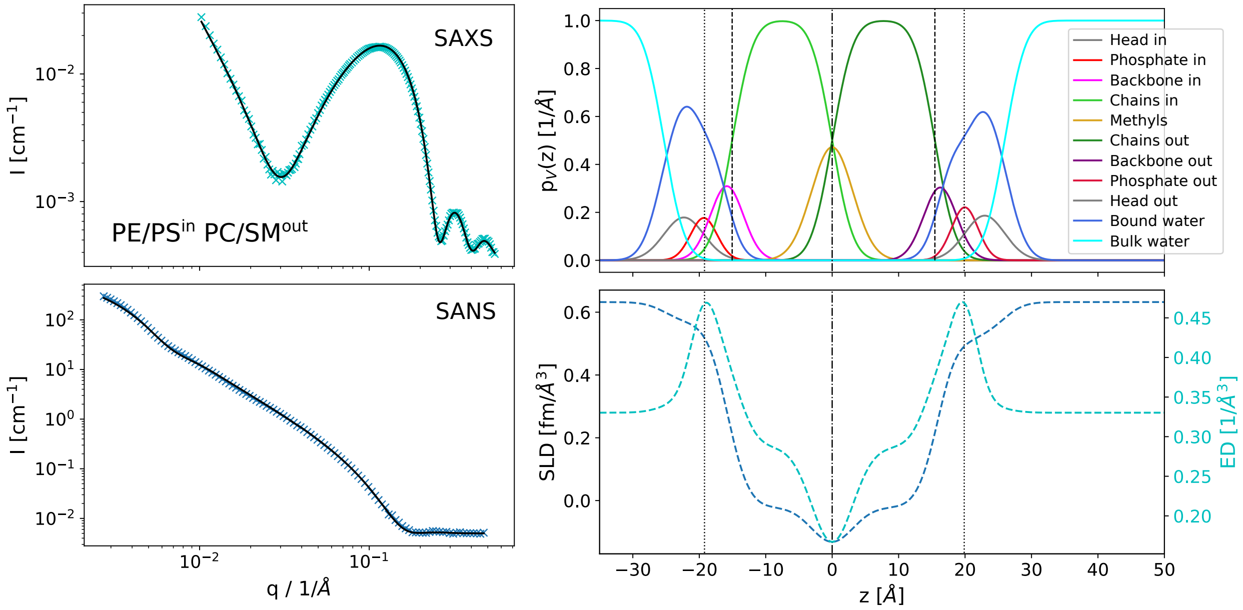


Figure S9: SAXS and SANS data with fits (black lines); SDP volume probability, electron density and neutron scattering length density profiles for the system $(PE/PS)^{in}/(PC/SM)^{out}$.

### 7 SAS-model parameters of reference LUVs

Table S5: SAS-fitting parameters of symmetric reference LUVs for POPE/POPG acceptor vesicles.
Inner leaflet samples (in): 90% POPE/POPG (9:1 mol/mol) and 10% POPC, MSM, or POPC/MSM (1:1 mol/mol).
Outer leaflet samples (out): 30% POPE/POPG (9:1 mol/mol) and 70% POPC, MSM, or POPC/MSM (1:1 mol/mol).
Scrambled vesicles (scram): same composition as aLUVs, see Tab. S4.

| | | POPC | | | MSM | | | PC/SM | | |
|---|---|---|---|---|---|---|---|---|---|---|
| | $\epsilon$ [%] | in | out | scram | in | out | scram | in | out | scram |
| $V_L^*$ [Å$^3$] | | 1202.1 | 1252.0 | 1212.94 | 1208.1 | 1293.56 | 1240.85 | 1205.11 | 1272.77 | 1247.96 |
| $V_H^*$ [Å$^3$] | | 257.4 | 304.48 | 267.63 | 252.0 | 266.68 | 257.65 | 254.74 | 285.58 | 274.27 |
| $r_{BB}^*$ | | 0.50 | 0.46 | 0.49 | 0.49 | 0.38 | 0.45 | 0.5 | 0.42 | 0.45 |
| $r_P^*$ | | 0.16 | 0.25 | 0.18 | 0.16 | 0.27 | 0.2 | 0.16 | 0.26 | 0.22 |
| $r^*$ | | 2.09 | 2.09 | 2.09 | 2.09 | 2 | 2.09 | 2.09 | 2.09 | 2.09 |
| $r_{12}^*$ | | 0.8 | 0.8 | 0.8 | 0.8 | 1 | 0.8 | 0.8 | 0.8 | 0.8 |
| $D_B$ [Å] | 3 | 39.6 | 38.9 | 37.37 | 40.6 | 43.13 | 38.87 | 39.92 | 41.28 | 38.36 |
| $D_{HH}$ [Å] | 3 | 37.2 | 35.6 | 33.84 | 37.0 | 39.44 | 35.59 | 36.3 | 37.53 | 35.31 |
| $2D_C$ [Å] | 3 | 31.1 | 29.4 | 29.12 | 32.2 | 34.23 | 30.79 | 31.47 | 32.01 | 29.92 |
| $D_{H1}$ [Å] | 20 | 3.1 | 3.1 | 2.36 | 2.4 | 2.61 | 2.4 | 2.41 | 2.76 | 2.7 |
| $A$ [Å$^2$] | 2 | 60.8 | 64.5 | 64.93 | 59.5 | 60 | 63.86 | 60.39 | 61.68 | 65.08 |
| $z_{BB}$ [Å] | 8 | 17.9 | 17.1 | 15.65 | 17.7 | 18.24 | 16.53 | 17.13 | 18.04 | 16.63 |
| $\sigma_{BB}$ [Å] | 20 | 2.5 | 2.5 | 2.5 | 2.5 | 2.5 | 2.5 | 2.5 | 2.5 | 2.5 |
| $z_P$ [Å] | 8 | 19.1 | 18.2 | 17.66 | 18.9 | 20.83 | 18.81 | 19.15 | 19.21 | 18.21 |
| $\sigma_P$ [Å] | 20 | 3.0 | 3.2 | 3.2 | 3.0 | 3 | 3.3 | 3.5 | 3.01 | 3.05 |
| $z_h$ [Å] | 3 | 22.0 | 21.5 | 19.95 | 23 | 23.72 | 21.37 | 23.06 | 21.1 | 20.02 |
| $\sigma_h^\dagger$ [Å] | | 2.8 | 2.8 | 2.8 | 2.8 | 2.8 | 2.8 | 2.8 | 2.8 | 2.8 |
| $\sigma_{HC}$ [Å] | | 2.5 | 2.5 | 2.5 | 2.5 | 2.5 | 2.5 | 2.5 | 2.5 | 2.5 |
| $\sigma_T$ [Å] | 5 | 2.4 | 2.4 | 2.79 | 2.4 | 4 | 3.76 | 3.51 | 2.7 | 4 |
| $\sigma_{poly}$ [%] | 6 | 4.8 | 3.22 | 5.96 | 5.4 | 6.34 | 4.82 | 4.31 | 3.66 | 7.5 |
| $V_{W,bound}$ [Å$^3$] | 6 | 29.8 | 29.9 | 28.83 | 29.5 | 29.8 | 29.01 | 29.75 | 29.87 | 28.47 |
| $n_W$ | 6 | 10.0 | 10.3 | 8.73 | 10.8 | 9.83 | 10 | 11.77 | 6.66 | 7.84 |
| $R_m$ [Å] | 10 | 370.1 | 370.2 | 430.7 | 372.5 | 470.2 | 408.6 | 356.2 | 391.4 | 390.6 |
| $\sigma_R$ [Å] | 10 | 103.9 | 123.7 | 171.7 | 102.3 | 127.7 | 182.9 | 91.67 | 106.7 | 188.3 |

Table S6: SAS-fitting parameters of symmetric reference LUVs for POPE/POPS acceptor vesicles.
Inner leaflet samples (in): 90% POPE/POPS (7:3 mol/mol) and 10% POPC, MSM, or POPC/MSM (1:1 mol/mol).
Outer leaflet samples (out): 30% POPE/POPS (7:3 mol/mol) and 70% POPC, MSM, or POPC/MSM (1:1 mol/mol).
Scrambled vesicles (scram): same composition as aLUVs, see Tab. S4.

| | | POPC | | | MSM | | | PC/SM | | |
|---|---|---|---|---|---|---|---|---|---|---|
| | $\epsilon$ [%] | in | out | scram | in | out | scram | in | out | scram |
| $V_L^*$ [Å$^3$] | | 1207.1 | 1253.6 | 1221.8 | 1213.03 | 1295.21 | 1255.49 | 1210.06 | 1274.42 | 1240.09 |
| $V_H^*$ [Å$^3$] | | 262.2 | 306.07 | 276.1 | 256.81 | 268.27 | 262.73 | 259.51 | 287.17 | 272.42 |
| $r_{BB}^*$ | | 0.49 | 0.46 | 0.48 | 0.48 | 0.37 | 0.42 | 0.48 | 0.41 | 0.45 |
| $r_P^*$ | | 0.19 | 0.26 | 0.21 | 0.19 | 0.28 | 0.24 | 0.19 | 0.27 | 0.23 |
| $r^*$ | | 2.09 | 2.09 | 2.09 | 2.09 | 2 | 2.09 | 2.09 | 2.09 | 2.09 |
| $r_{12}^*$ | | 0.8 | 0.8 | 0.8 | 0.8 | 1 | 0.8 | 0.8 | 0.8 | 0.8 |
| $D_B$ [Å] | 3 | 37.8 | 38.2 | 37.2 | 38.22 | 42.65 | 39.08 | 38.06 | 40.22 | 37.09 |
| $D_{HH}$ [Å] | 3 | 37.0 | 35.3 | 32.7 | 36.85 | 38.99 | 38.93 | 36.28 | 37.2 | 36.53 |
| $2D_C$ [Å] | 3 | 29.6 | 28.8 | 28.8 | 30.12 | 33.81 | 30.89 | 29.89 | 31.15 | 28.93 |
| $D_{H1}$ [Å] | 20 | 3.7 | 3.2 | 2.0 | 3.37 | 2.59 | 4.02 | 3.19 | 3.03 | 3.8 |
| $A$ [Å$^2$] | 2 | 63.8 | 65.7 | 65.8 | 63.49 | 60.75 | 64.27 | 63.61 | 63.39 | 66.89 |
| $z_{BB}$ [Å] | 8 | 16.3 | 16.1 | 16.0 | 17.22 | 18.53 | 19.39 | 16.8 | 17.84 | 17.31 |
| $\sigma_{BB}$ [Å] | 20 | 2.5 | 2.5 | 2.5 | 2.5 | 2.5 | 2.86 | 2.5 | 2.5 | 2.5 |
| $z_P$ [Å] | 8 | 19.1 | 18.3 | 16.0 | 19.92 | 20.34 | 19.39 | 19.59 | 19.1 | 19.42 |
| $\sigma_P$ [Å] | 20 | 2.2 | 2.8 | 2.9 | 3.5 | 3.29 | 2.2 | 3.5 | 3.27 | 3.5 |
| $z_h$ [Å] | 3 | 24.1 | 23.3 | 21 | 19.92 | 21.77 | 24.39 | 19.59 | 19.47 | 19.42 |
| $\sigma_h^\dagger$ [Å] | | 2.8 | 2.8 | 2.8 | 2.8 | 2.8 | 2.8 | 2.8 | 2.8 | 2.8 |
| $\sigma_{HC}$ [Å] | | 2.5 | 2.5 | 2.5 | 2.5 | 2.5 | 2.5 | 2.5 | 2.5 | 2.5 |
| $\sigma_T$ [Å] | 5 | 3.5 | 3.1 | 4.0 | 3.32 | 3.89 | 2.4 | 3.26 | 3.19 | 4 |
| $\sigma_{poly}$ [%] | 6 | 7.3 | 6.63 | 10.0 | 5.26 | 4.81 | 9.46 | 6.08 | 5.58 | 6.92 |
| $V_{W,bound}$ [Å$^3$] | 6 | 30.0 | 29.7 | 30.3 | 30.03 | 29.78 | 30.01 | 28.97 | 29.26 | 29.41 |
| $n_W$ | 6 | 16.9 | 15.2 | 11.4 | 7.58 | 6.53 | 16.25 | 7.07 | 4.54 | 8.14 |
| $R_m$ [Å] | 10 | 381 | 392.6 | 469.1 | 409.5 | 495.7 | 607.7 | 430.9 | 451.9 | 500.7 |
| $\sigma_R$ [Å] | 10 | 109.2 | 133.5 | 154.4 | 131.7 | 167.1 | 151.9 | 140.6 | 133.1 | 128.7 |

Table S7: SAS-fitting parameters of symmetric reference LUVs for POPE/POPG acceptor vesicles.
Inner leaflet samples (in): 90% POPE/POPG (9:1 mol/mol) and 10% ESM.
Outer leaflet samples (out): 30% POPE/POPG (9:1 mol/mol) and 70% ESM.
Scrambled vesicles (scram): same composition as aLUVs, see Tab. S4.

| | $\epsilon$ [%] | ESM in | out | scram |
|---|---|---|---|---|
| $V_L^*$ [Å$^3$] | | 1196.5 | 1212.2 | 1181.0 |
| $V_H^*$ [Å$^3$] | | 252.0 | 266.68 | 259.1 |
| $r_{BB}^*$ | | 0.49 | 0.38 | 0.44 |
| $r_P^*$ | | 0.16 | 0.27 | 0.21 |
| $r^*$ | | 2.09 | 2.09 | 2.09 |
| $r_{12}^*$ | | 0.8 | 0.8 | 0.8 |
| $D_B$ [Å] | 3 | 38.0 | 41.1 | 38.2 |
| $D_{HH}$ [Å] | 3 | 36.9 | 37.0 | 34.9 |
| $2D_C$ [Å] | 3 | 30.0 | 32.0 | 29.8 |
| $D_{H1}$ [Å] | 20 | 3.4 | 2.5 | 2.6 |
| $A$ [Å$^2$] | 2 | 63.0 | 59.1 | 61.8 |
| $z_{BB}$ [Å] | 8 | 17.6 | 17.6 | 15.7 |
| $\sigma_{BB}$ [Å] | 20 | 2.5 | 2.5 | 2.5 |
| $z_P$ [Å] | 8 | 19.4 | 18.7 | 20.2 |
| $\sigma_P$ [Å] | 20 | 3.5 | 2.7 | 3.5 |
| $z_h$ [Å] | 3 | 19.4 | 22.9 | 21 |
| $\sigma_h^\dagger$ [Å] | | 2.8 | 2.8 | 2.8 |
| $\sigma_{HC}$ [Å] | | 2.5 | 2.5 | 2.5 |
| $\sigma_T$ [Å] | 5 | 3.1 | 2.4 | 4.0 |
| $\sigma_{poly}$ [%] | 6 | 5.3 | 4.95 | 8.5 |
| $V_{W,bound}$ [Å$^3$] | 6 | 27.7 | 27.6 | 30.3 |
| $n_W$ | 6 | 6.7 | 10.0 | 8.8 |
| $R_m$ [Å] | 10 | 393.1 | 455 | 531.6 |
| $\sigma_R$ [Å] | 10 | 114.1 | 132.9 | 138.9 |

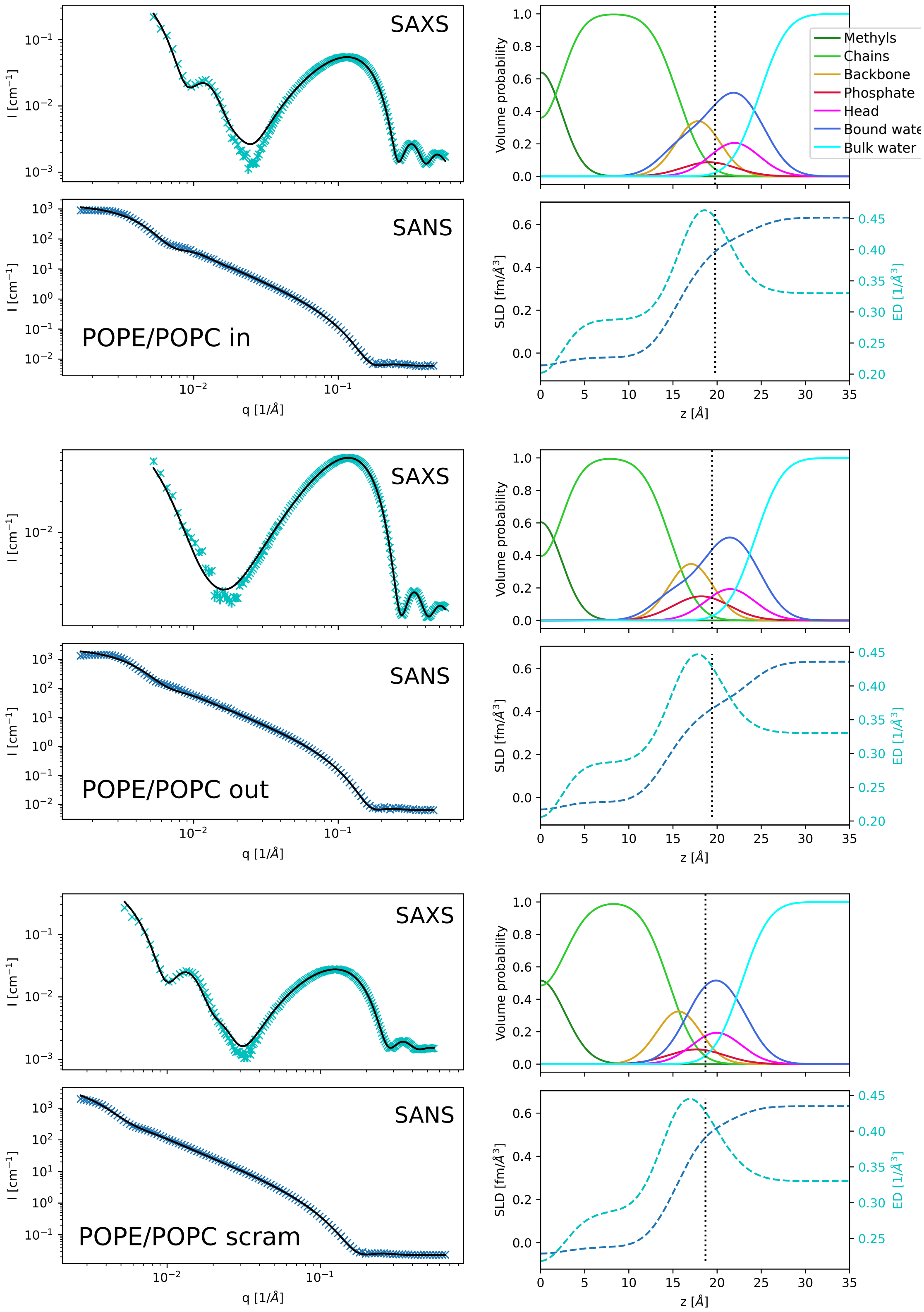


Figure S10: SAXS and SANS data with fits (black lines); SDP volume probability, electron density and neutron scattering length density profiles for POPE/POPC inner/outer leaflet symmetric mimics, as well as the scrambled sample.

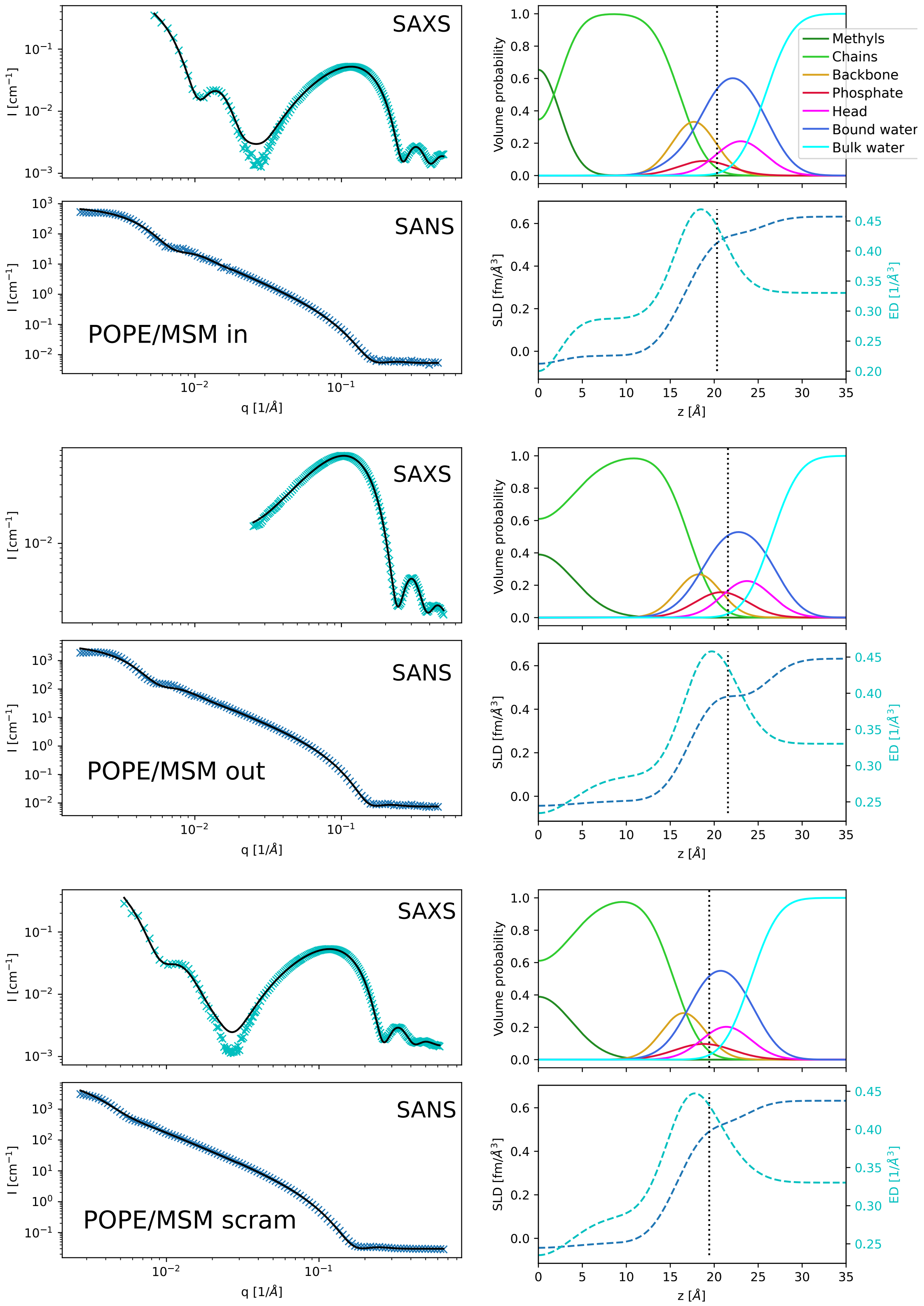


Figure S11: SAXS and SANS data with fits (black lines); SDP volume probability, electron density and neutron scattering length density profiles for POPE/MSM inner/outer leaflet symmetric mimics, as well as the scrambled sample.

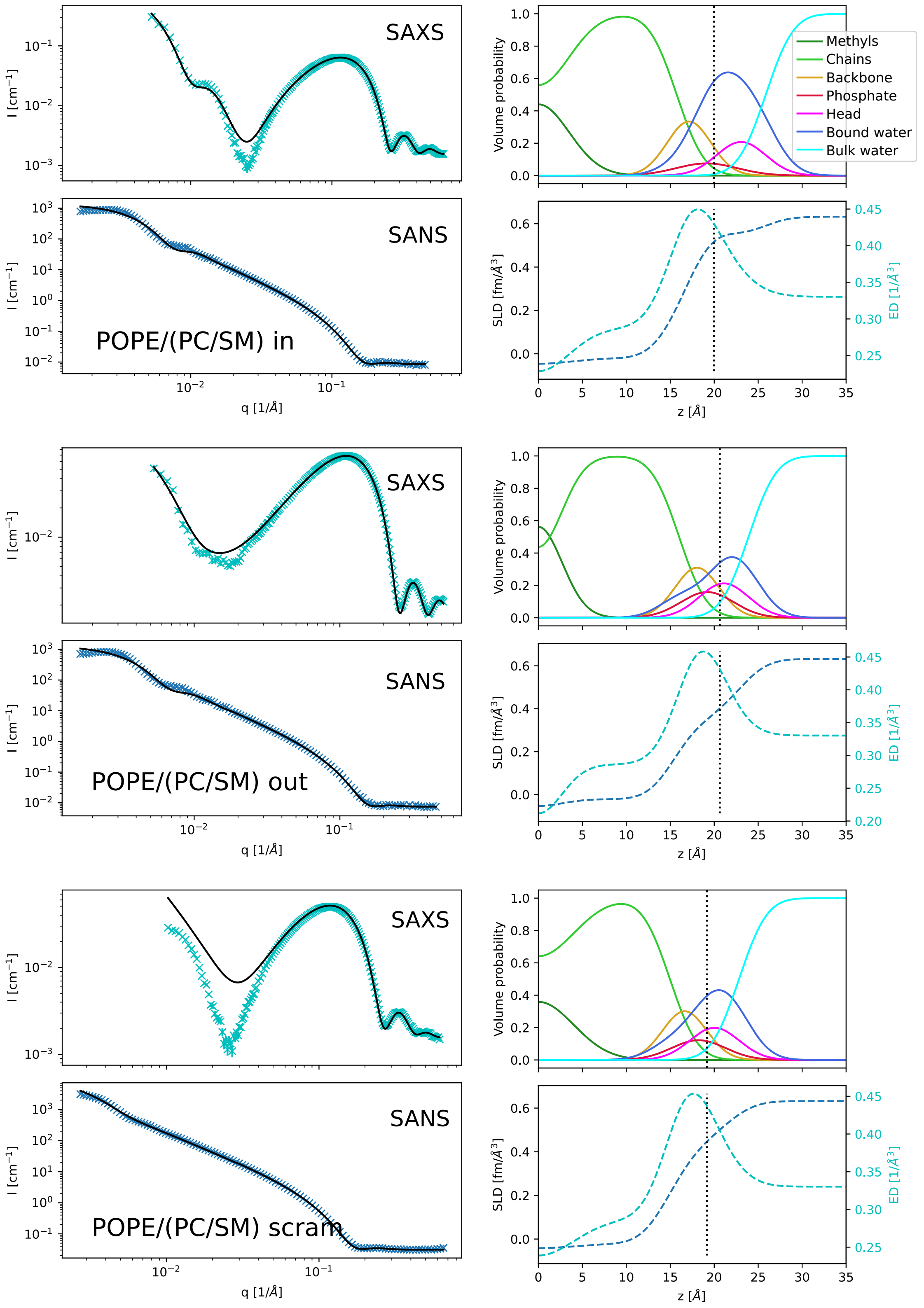


Figure S12: SAXS and SANS data with fits (black lines); SDP volume probability, electron density and neutron scattering length density profiles for POPE/(PC/SM) inner/outer leaflet symmetric mimics, as well as the scrambled sample.

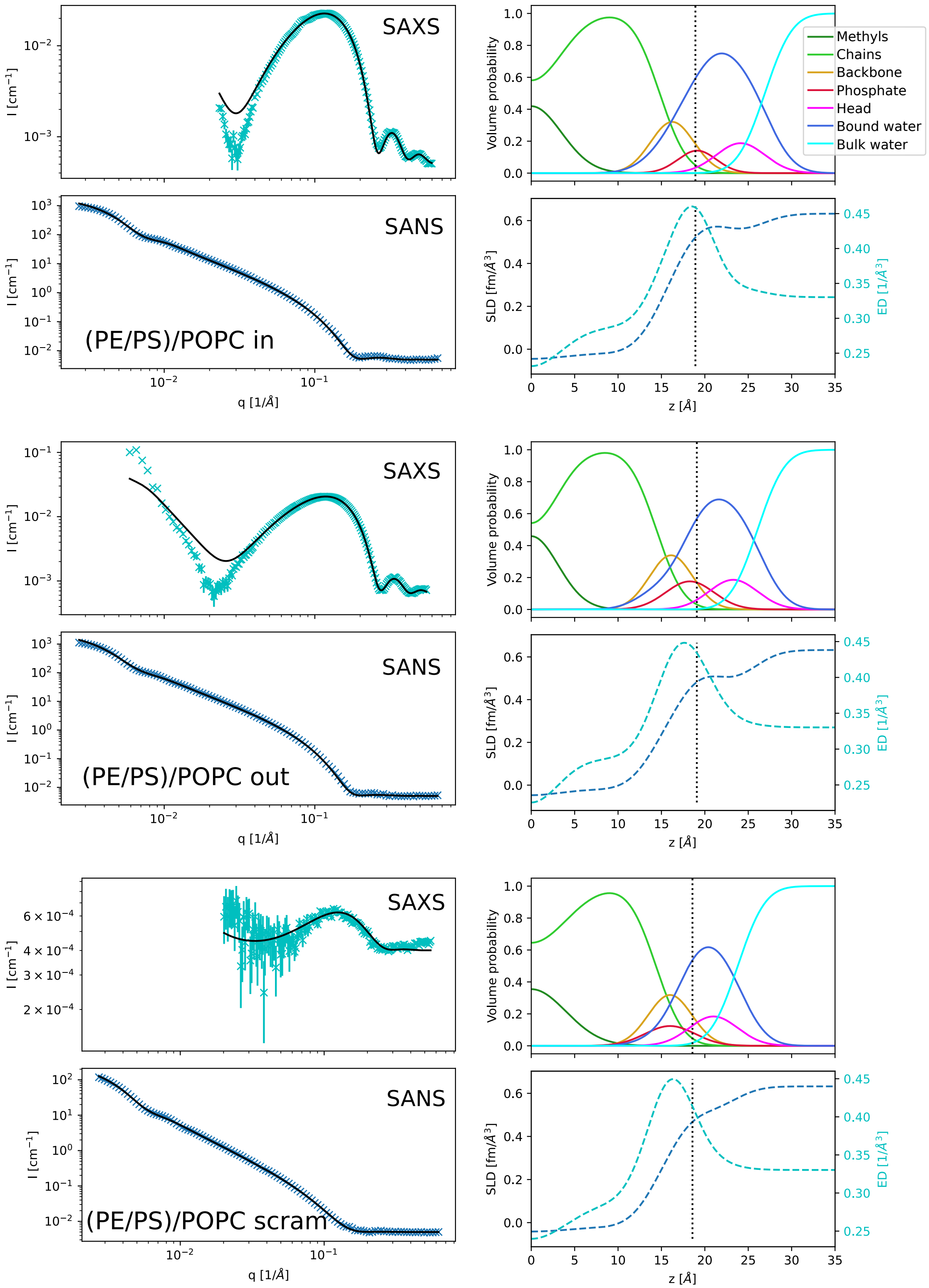


Figure S13: SAXS and SANS data with fits (black lines); SDP volume probability, electron density and neutron scattering length density profiles for (PE/PS)/POPC inner/outer leaflet symmetric mimics, as well as the scrambled sample.

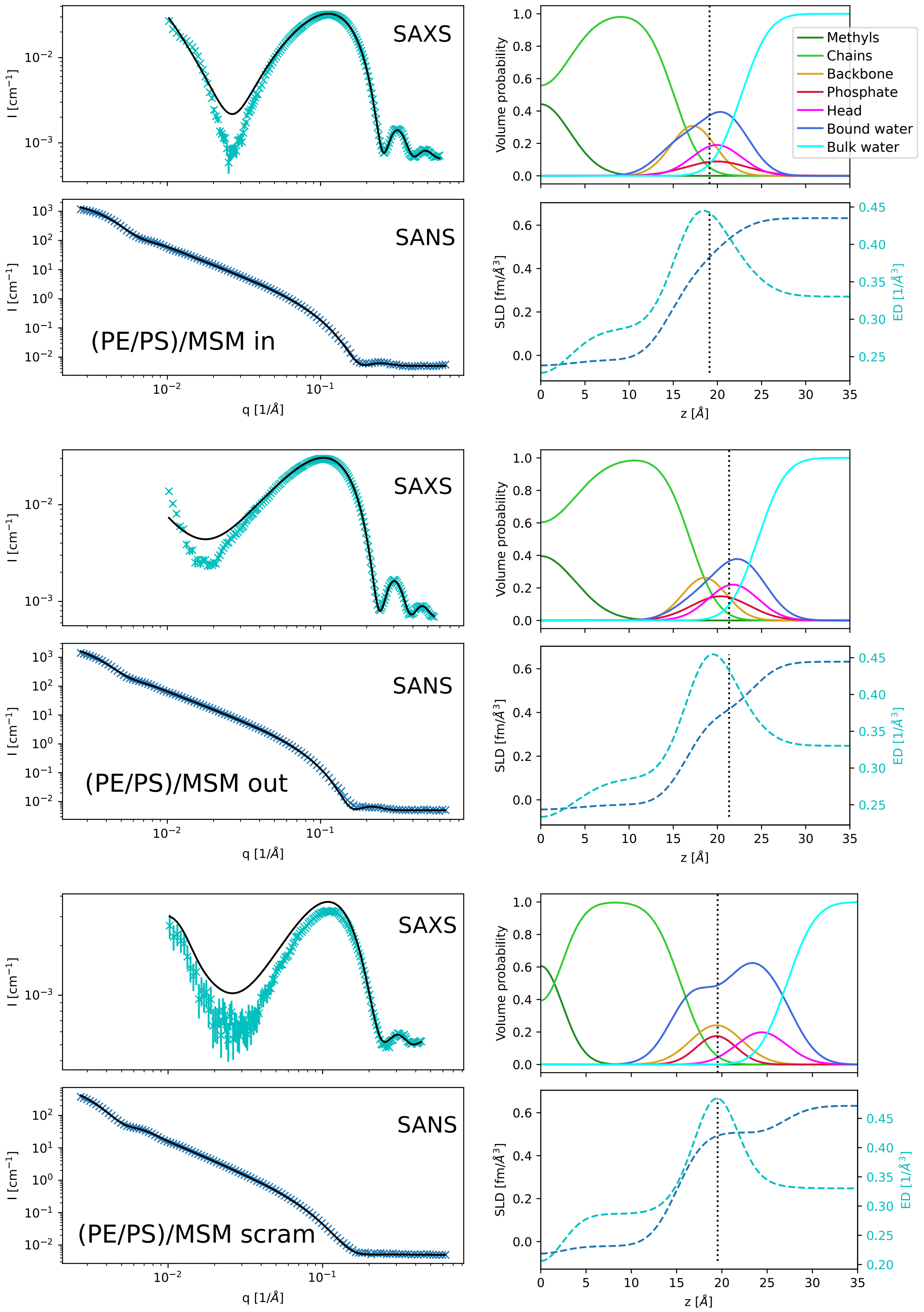


Figure S14: SAXS and SANS data with fits (black lines); SDP volume probability, electron density and neutron scattering length density profiles for (PE/PS)/MSM inner/outer leaflet symmetric mimics, as well as the scrambled sample.

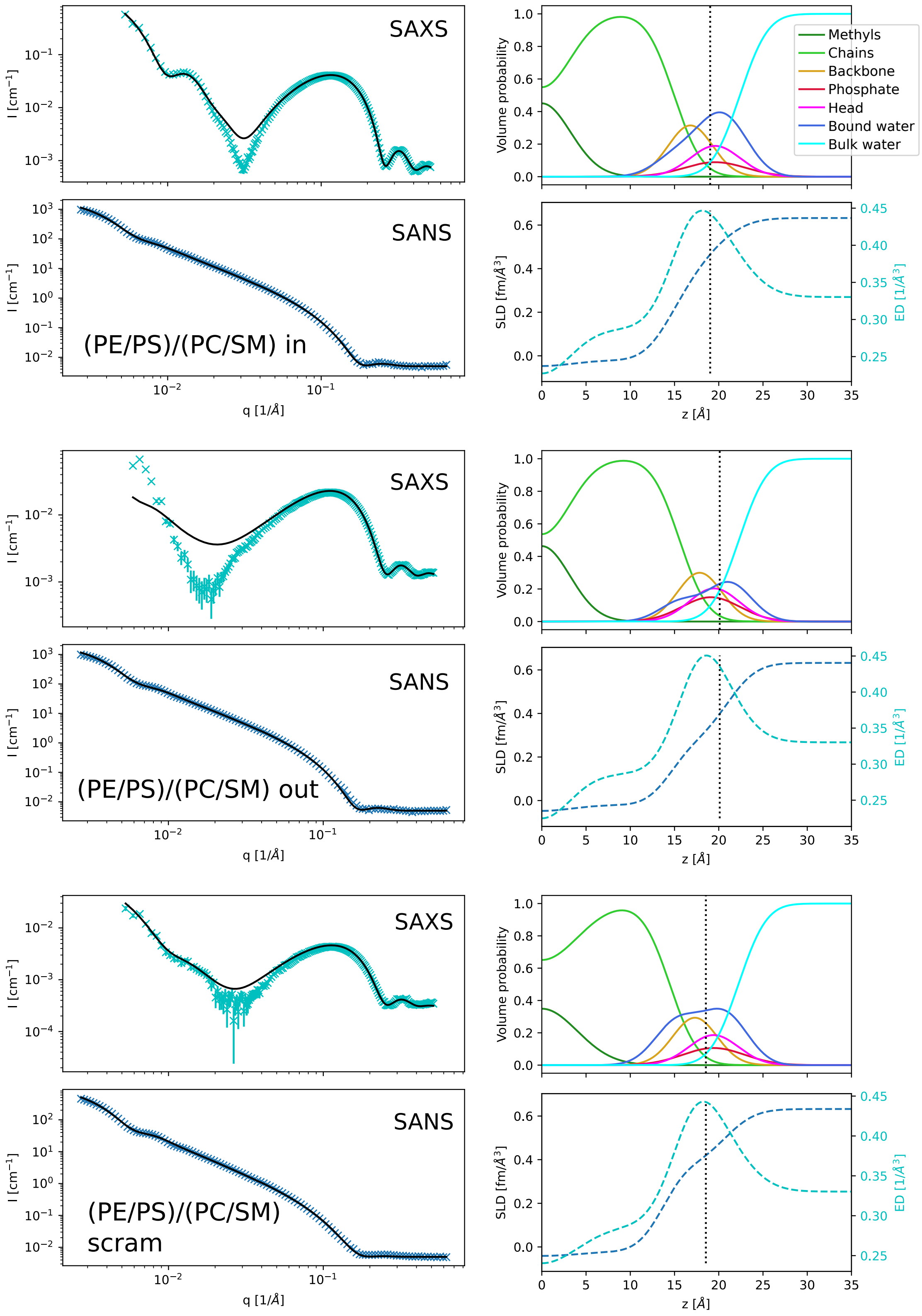


Figure S15: SAXS and SANS data with fits (black lines); SDP volume probability, electron density and neutron scattering length density profiles for (PE/PS)/(PC/SM) inner/outer leaflet symmetric mimics, as well as the scrambled sample.

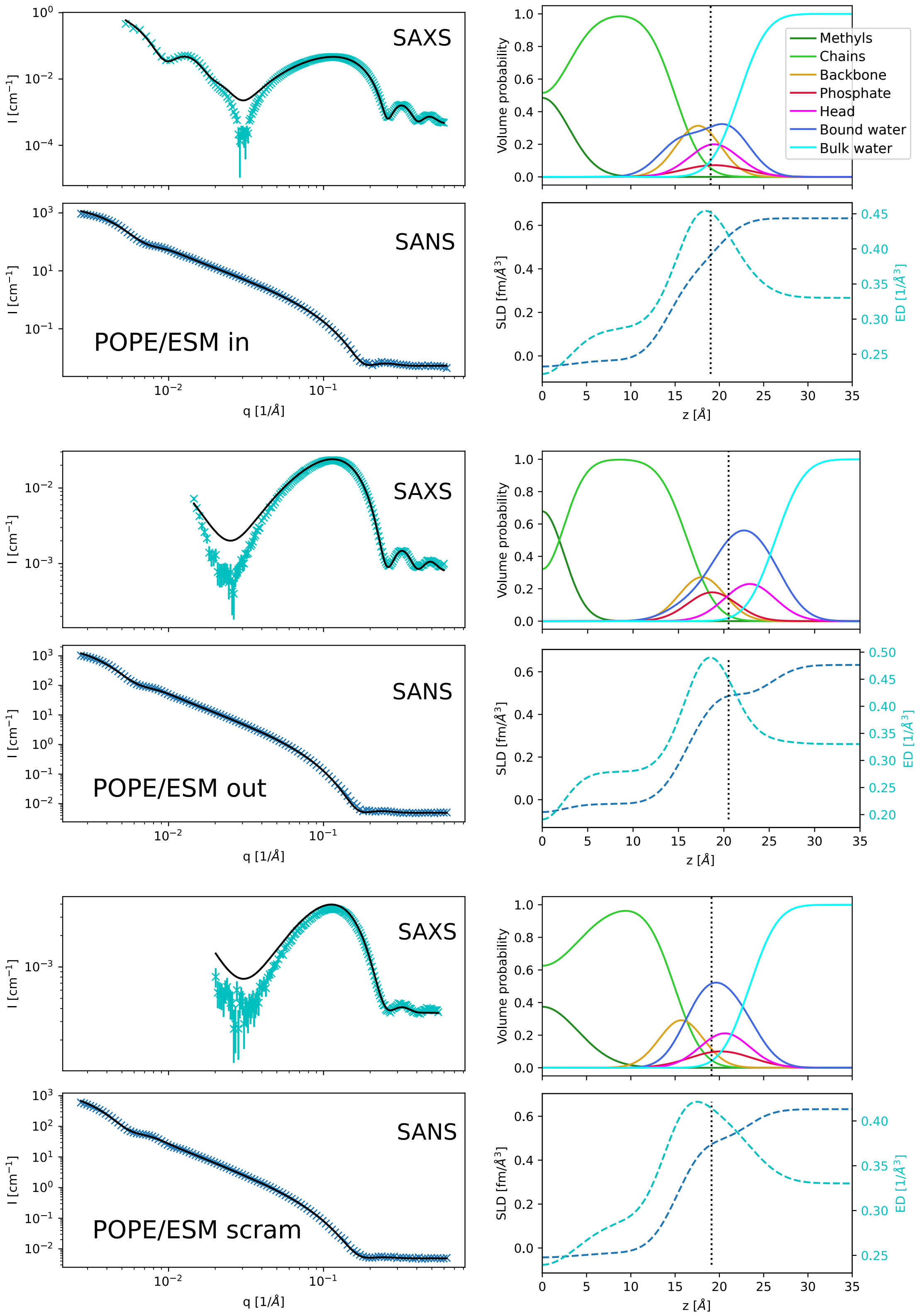


Figure S16: SAXS and SANS data with fits (black lines); SDP volume probability, electron density and neutron scattering length density profiles for POPE/ESM inner/outer leaflet symmetric mimics, as well as the scrambled sample.
